# High Resolution Kinetic Characterization and Dynamic Mathematical Modeling of the RIG-I Signaling Pathway and the Antiviral Responses

**DOI:** 10.1101/2022.08.05.502818

**Authors:** Sandy S. Burkart, Darius Schweinoch, Jamie Frankish, Carola Sparn, Sandra Wüst, Christian Urban, Antonio Piras, Andreas Pichlmair, Joschka Willemsen, Lars Kaderali, Marco Binder

## Abstract

The pattern recognition receptor RIG-I is essential for the recognition of viral dsRNA and the activation of a cell-autonomous antiviral response. Upon stimulation, RIG-I triggers a signaling cascade leading to the expression of cytokines, most prominently type I and III interferons (IFNs). IFNs are secreted and signal in an auto- and paracrine manner to trigger the expression of a large variety of IFN-stimulated genes, which in concert establish an antiviral state of the cell. While the topology of this pathway has been studied quite intensively, the dynamics, particularly of the RIG-I-mediated IFN induction, is much less understood. Here, we employed electroporation-based transfection to synchronously activate the RIG-I signaling pathway, enabling us to characterize the kinetics and dynamics of cell-intrinsic innate immune signaling to virus infections. By employing an A549 IFNAR1/IFNLR deficient cell line, we could analyze the difference between the primary RIG-I signaling phase and the secondary signaling phase downstream of the IFN receptors. We further used our quantitative data to set up and calibrate a comprehensive dynamic mathematical model of the RIG-I and IFN signaling pathways. This model accurately predicts the kinetics of signaling events downstream of dsRNA recognition by RIG-I as well as the feedback and signal amplification by secreted IFN and JAK/STAT signaling. We have furthermore investigated the impact of various viral immune antagonists on the signaling dynamics experimentally, and we utilized the here described modelling approach to simulate and *in silico* study these critical virus-host interactions. Our work provides a comprehensive insight into the signaling events occurring early upon virus infection and opens up new avenues to study and disentangle the complexity of the host-virus interface.

## INTRODUCTION

Recognition of pathogen-associated molecular patterns (PAMPs) by a variety of cell-surface and intracellular pattern recognition receptors (PRRs), including toll-like receptors (TLRs) and retinoic acid-inducible gene-I-like receptors (RLRs), can trigger signaling cascades leading to the expression of cytokines e.g., interferons (IFNs), and interferon-stimulated genes (ISGs). Briefly, upon viral double-stranded RNA (dsRNA) recognition, retinoic acid-inducible gene-I (RIG-I) initiates a signaling cascade through mitochondrial antiviral-signaling protein (MAVS), leading to the activation and nuclear translocation of transcription factors such as IFN-regulatory factor 3 (IRF3) and nuclear factor kappa B (NF-κB). The activation of these transcription factors results in the production and secretion of mostly type I and III IFNs. Auto- and paracrine IFN signaling through the corresponding IFN receptors (IFNR) can further induce multiple downstream signaling pathways. The most important signaling axis is the JAK/STAT pathway, leading to the formation of a transcription factor composed of the signal transducer and activator of transcription 1 (STAT1), STAT2, and IRF9, in conjunction known as the IFN-stimulated gene factor 3 (ISGF3) complex. Upon activation, this complex binds to IFN-stimulated response elements (ISREs) within gene promoters, further leading to the induction of a large variety of ISGs, which exert numerous antiviral functions [1,2]. In addition, upon activation by the only type II IFN, known as IFN-γ, STAT1 can form homodimers that bind to IFN-gamma-activated-sequence (GAS) elements in promoter regions of ISGs. Thus, the expression of both, IFNs and ISGs, ultimately establishes an antiviral state of a cell [3–6]. In light of the important, often even decisive impact of the IFN system on the outcome of viral infection, viruses have evolved powerful strategies to evade innate immune recognition and defense [7,8]. One strategy is to omit recognition by PRRs in the first place, with hepatitis B virus being a prime example (“stealth virus”) [9,10]. However, this appears excessively difficult to achieve, and most viruses have rather evolved antagonists of host-cellular antiviral pathways (“cunning virus”) [10,11]. These virus-encoded factors have an ample choice of points of attack, and different virus groups and families target different cellular processes [12]. For instance, the classical swine fever virus (CSFV) protein N^pro^ was identified as an antagonist of RIG-I signaling by targeting IRF3 for degradation [13,14]. Similarly, beyond being indispensable for viral replication, the hepatitis C virus (HCV) protease NS3/4A additionally targets MAVS for cleavage and release from the mitochondrial membrane, rendering RLR/IRF3-signaling dysfunctional [15–17]. Viral antagonists have been studied extensively, and many have been characterized in detail. Nonetheless, most studies investigating such viral antagonists are limited to robust overexpression and end-point determination of the degree of inhibition of the host response. However, in an actual infection, active concentrations of the antagonists are gradually increasing as the virus replicates and expresses its genes. Moreover, depending on the very target of inhibition, the resulting effect may either delay the mounting of an antiviral response or lead to an overall lower amplitude of the response (reviewed in [7,8,18]). Therefore, it is very important to investigate the dynamics of the IFN response to determine the outcome of an infection as well as the dynamic impact of antagonists to ultimately understand viral immune evasion. Given this outstanding importance of the above-mentioned dynamics (i.e., combination of kinetics and magnitude), it is essential to gain a profound understanding of the cellular antiviral signaling system. Particularly for viruses, this system is based largely on cytoplasmic sensors, such as the RIG-I-like receptors. While simultaneous activation of cell surface receptors such as TNF receptor [19], IL-1 receptor [20], and some toll-like receptors [21] can easily be achieved by ligand addition to the cell culture media, this is particularly challenging for intracellular receptors such as RIG-I. Owing to this cytosolic localization of the receptors, instant stimulation of the pathway is challenging, hence virtually all kinetic studies were based on actual virus infection or liposome-based transfection of virus-like RNA [22,23]. However, cellular uptake of the PAMP (i.e., viral RNA) in those cases depends on endocytic processes, which introduce a large variability in the timing of the PAMP to be accessible to PRRs.

In this study, we kinetically characterized the dynamics of antiviral signaling triggered by RIG-I in human alveolar epithelial (A549) cells and established a mathematical model, which allows accurate dynamic simulation. We established an approach permitting the synchronous stimulation of A549 cells with virus-like dsRNA, allowing for a finer resolution of pathway kinetics. With this system, we analyzed phosphorylation and expression of critical proteins in the RIG-I signaling cascade in a fine-grained time course upon synchronous stimulation using live-cell imaging, quantitative western blotting, and qRT-PCR. We then used these time-resolved data to establish and calibrate a mathematical model that reproduces the kinetics of key signaling events within the core RIG-I pathway. In a next step, we combined our RIG-I core model with a previously published and validated model of the IFN signaling [24], ultimately generating a comprehensive model of the full innate antiviral system. Although the published model was trained on kinetic measurements in a different cell line, we could employ all reaction rates downstream of the IFN receptor from the original model without re-adjustments, which corroborates the high conservation of antiviral signaling across different cell lines. Furthermore, using viral antagonists with well-established mechanisms of action, we characterized their impact on RIG-I and IFN signaling dynamics and compared experimental data to model simulations. Previous models combining RIG-I and IFN pathway components have been limited by the availability of experimental data [25,26], were of reduced complexity compared to the herein presented model (e.g., [27]), or focused on RIG-I pathway components [28] (eminently reviewed in [29]). We present the most comprehensive, data-based mathematical model of the cell-intrinsic antiviral defense system, permitting simulation and analysis of critical virus-host interactions early into infection.

## RESULTS

### RIG-I signaling upon dsRNA recognition is highly deterministic and synchronous

In previous studies, RLR signaling kinetics upon viral infection or liposome-based transfection resulted in a heterogeneous nuclear translocation of transcription factors, such as IRF3 or IRF7, or expression of the IFNB1 gene [23,30]. For instance, Rand and colleagues reported a cell-intrinsic stochasticity in the activation of IRF7 and NF-κB upon virus infection of murine cells [23]. To determine the dynamics of RLR signaling, we employed A549 cells stably co-expressing cytosolic IRF3-eGFP and nuclear H2B-mCherry and inspected nuclear translocation of IRF3 after RIG-I stimulation (Figure 1). As a ligand highly specific for RIG-I, 400 bp 5’ppp*-*dsRNA [31] was delivered to cells either by liposome- or by electroporation-based transfection. In contrast to literature, we found RIG-I signaling to be highly deterministic and synchronous after 5’ppp*-*dsRNA delivery by electroporation-based transfection, while mock electro-transfection did not induce innate immune signaling (Supplementary Figure S1). This suggests that the previously observed stochasticity was largely due to staggered uptake of the stimulatory RNA during infection. Confocal microscopy analysis of IRF3-eGFP nuclear translocation kinetics (Figure 1A) demonstrated a clear difference between the methods of RLR ligand delivery. While electro-transfected cells showed a rapid and synchronous IRF3 translocation, liposome-based transfection of 5’ppp-dsRNA resulted in a delayed and overall slower translocation in A549 cells (Figure 1A). In order to quantitatively assess nuclear translocation of IRF3-eGFP upon those different stimulation approaches, we analyzed co-localization of IRF3-eGFP with H2B*-*mCherry over time using the Incucyte live-cell imaging system and, further, used the ilastik software for automated quantification of at least 500 single cells per condition and time point. Using live-cell imaging to quantitatively assess nuclear translocation after RIG-I stimulation, we again observed that liposome-based transfection of 5’ppp-dsRNA resulted in a constant increase of the fraction of cells exhibiting nuclear IRF3-eGFP over six hours, whereas electro-transfection induced a synchronous and rapid increase starting 30 minutes after stimulation (Figure 1B). To determine whether the observed results were due to possible differences in the RNA amounts successfully entering the cytosol, we titrated the amount of 5’ppp-dsRNA. While indeed activation kinetics were slightly affected and the maximum fraction of activated cells was significantly affected by decreasing dsRNA concentrations, the qualitative characteristics remained fully consistent: independent of the dsRNA amount, electro-transfection led to substantially steeper activation kinetics (Figure 1C) as compared to liposome-based transfection (Figure 1D). This corroborated our hypothesis of uptake modalities playing the single most decisive role in the previously described asynchronicity (“stochasticity”) of RLR-driven IRF responses.

**Figure 1.**
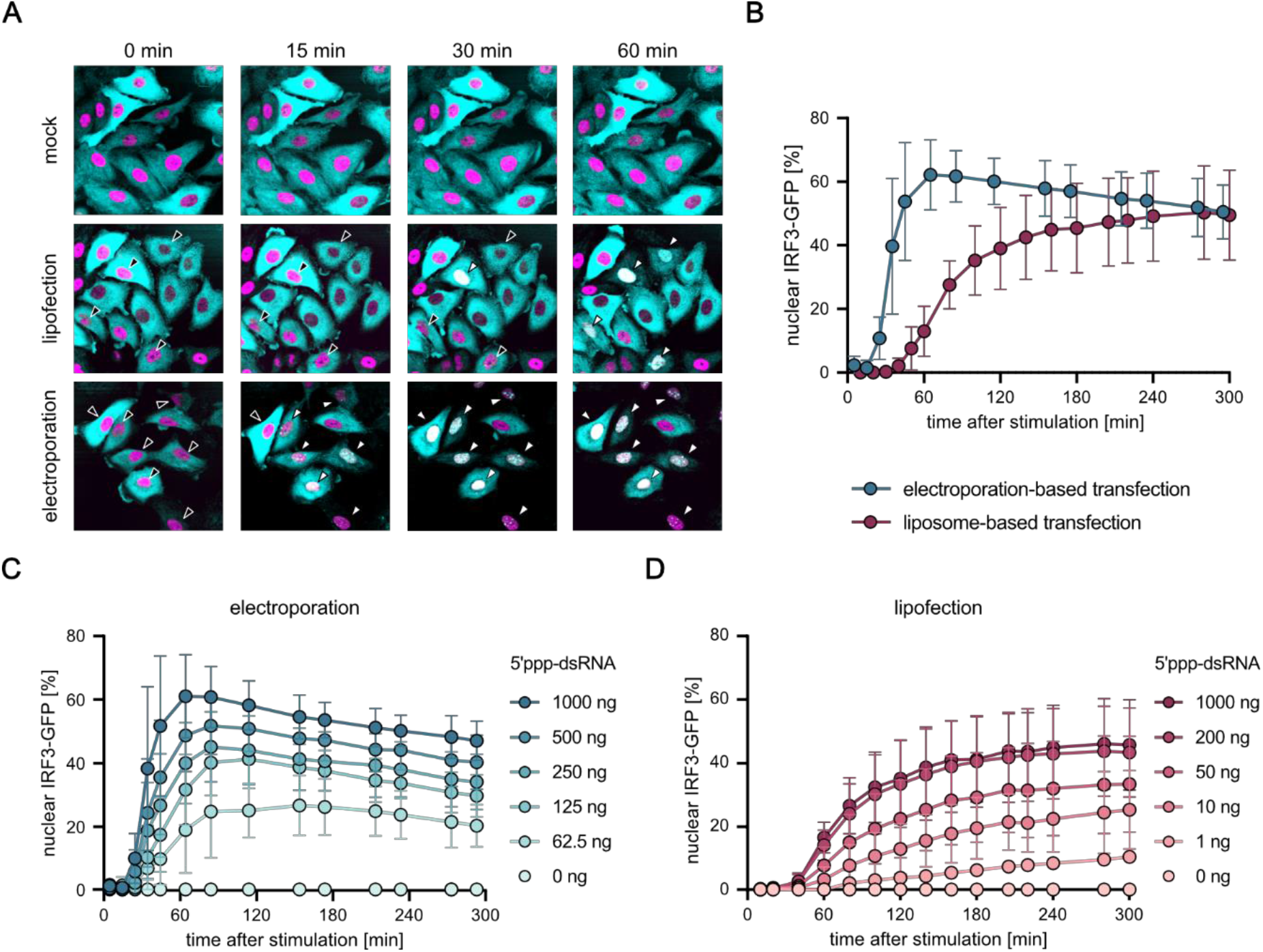
Synchronous activation of the RIG-I signaling pathway upon dsRNA electro-transfection. **(A)** Confocal microscopy time course of A549 cells stably expressing cytoplasmic IRF3-eGFP (cyan) and the nuclear marker H2B-mCherry (magenta) upon different stimulation approaches. Cells were either mock-treated or stimulated with 1 µg *in vitro* generated 5’ppp-dsRNA using a liposome-based transfection or in-well electro-transfection using the Lonza Nucleofector® system. Black arrowheads indicate cytoplasmic IRF3-eGFP, white arrowheads indicate translocated, nuclear IRF3-eGFP. **(B)** Quantification and comparison of IRF3-eGFP nuclear translocation upon stimulation with 1 µg 5’ppp-dsRNA using a liposome-based or electroporation-based transfection method. Nuclear translocation of IRF3-eGFP and co-localization with H2B-mCherry was analyzed over time using the Incucyte live*-*cell imaging system. Here, at least 500 and up to 2500 individual cells were automatically evaluated for each time point in each condition using the ilastik software. **(C)** Quantification and comparison of IRF3-eGFP nuclear translocation upon electroporation*-*based transfection of varying concentrations of 5’ppp-dsRNA. **(D)** Quantification and comparison of IRF3-eGFP nuclear translocation upon liposome-based transfection of varying concentrations of 5’ppp-dsRNA. Graphs depict representative images (A), mean ± SD of four (B) or three (C*-*D) biologically independent experiments, respectively.

### Synchronous RIG-I stimulation results in a fast onset of RLR pathway signaling

The highly synchronous activation of the RLR signaling pathway by electro-transfection permitted us to kinetically characterize the RIG-I signaling dynamics in more detail. We synchronously stimulated A549 wild type (wt) cells with 5’ppp-dsRNA and analyzed the activation (i.e., phosphorylation) status of RLR signaling pathway components or protein abundance (for IκB) within the IRF3- (Figure 2A) or NF-κB-axis (Figure 2B) over time via western blotting. Already ten minutes after stimulation, activated and phosphorylated forms of the kinases TBK1 and IKKε (for IRF3) or IKKβ (for NF-κB) were detectable.

**Figure 2.**
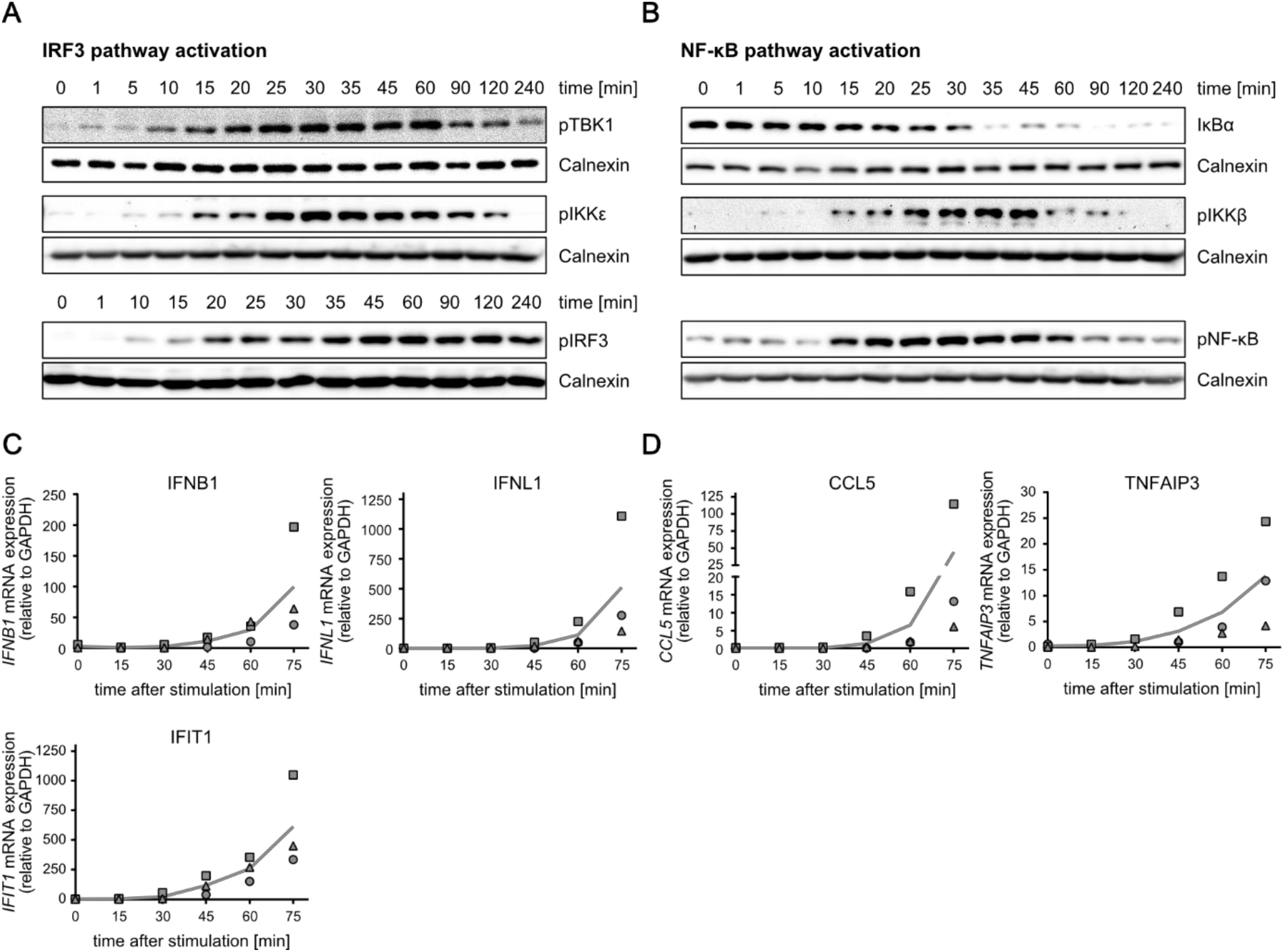
Time-resolved activation of the early RIG-I signaling cascade. Synchronized stimulation of A549 wt cells with 220 ng 5’ppp-dsRNA by electro-transfection was used to kinetically characterize the dynamics of the RIG-I signaling pathway. Protein abundance and phosphorylation status of RLR signaling pathway components within the **(A)** IRF3 or **(B)** NF-κB axis were determined using western blot analysis. Early mRNA expression onset of **(C)** IRF3-dependent genes (IFNB1, IFNL1, and IFIT1) and **(D)** NF-κB-dependent genes (CCL5 and TNFAIP3) after dsRNA electro-transfection was measured using quantitative RT-PCR. Values were normalized to the housekeeping gene GAPDH using the 2^-ΔCt^ method. Graphs depict representative blots (A, B) or mean and individual, biological replicate values (C, D) of three biologically independent experiments. In fact, the earlier onset of decay of the NF κB inhibitor, IκBα, five minutes after transfection, which is dependent on IKKβ activation, argues for an even earlier kinase activation (Figure 2B). IRF3 phosphorylation started shortly after TBK1 and IKKε activation (Figure 2A). Furthermore, while NF κB and IKKβ kinase phosphorylation decreased concurrently (Figure 2B), IRF3 phosphorylation was surprisingly stable within the experimental time frame, despite pTBK1 and pIKKε levels declining from one hour onwards (Figure 2A). Consistently with the rapid onset of RIG I signaling, early transcripts of the target genes IFNB1, IFNL1, and IFIT1 (Figure 2C) as well as CCL5 and TNFAIP3 (Figure 2D) were detectable already at 45 to 60 minutes post stimulation by qRT PCR.

### Dynamic RIG-I signaling model accurately reproduces activation kinetics of essential pathway components

Our acquired time-resolved, quantitative data on the activation kinetics of the canonical RIG-I pathway components provided an excellent description of the antiviral signaling dynamics in our experimental system. In order to generalize these observations and investigate kinetic characteristics such as rate-limiting steps in this important pathway, we employed kinetic mathematical modeling. We developed a computational model using ordinary differential equations (ODE) representing major steps within the RIG-I signaling pathway based on regular mass-action kinetics. Our model comprises 19 pathway components (“species”) distributed into two distinct compartments, the cytoplasm or the nucleus. We introduced 22 rate constants representing canonical steps of the RIG-I signaling cascade, as well as some additional parameters (e.g., normalization factors) to account for initial conditions, experimental conditions, and the relative nature of the quantitative immunoblot data (Figure 3A). In order to determine absolute concentrations of all relevant proteins, we acquired quantitative full-proteome data of naïve A549 cells by label-free mass spectrometry. Of the 22 kinetic rate constants (i.e., signal transduction and transport events), nine were fixed based on information from previous publications or initial conditions and the remaining 11 were optimized during model fitting (Supplementary Material). For model calibration, we used relative protein quantities and phosphorylation intensities from our quantitative immunoblot (Figure 3B) and transcript levels from qRT-PCR data (Figure 3C) of four biologically independent replicate experiments. Our model was able to accurately and reliably capture the measured signaling dynamics with only one unique set of parameters (Figure 2, Figure 3B, C, Supplementary Material).

**Figure 3.**
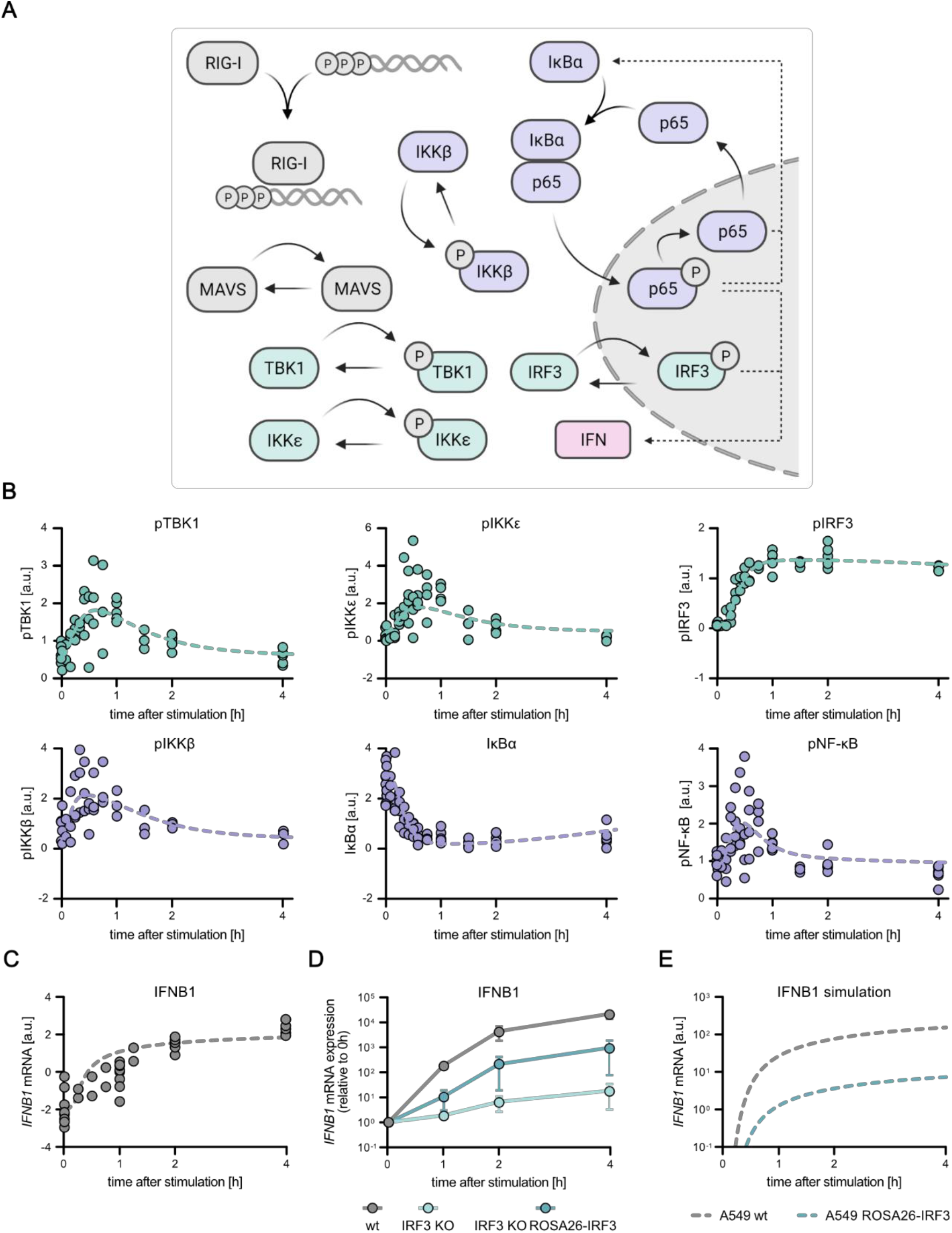
Mathematical model of the core RIG-I pathway based on quantitative data. The previously obtained, time-resolved data was used to establish and calibrate a mathematical model which reproduces the kinetics of key signaling events within the core RIG-I pathway. **(A)** Schematic depiction of key components relevant for the establishment of a mathematical model of the core RIG-I signaling pathway. **(B)** Quantitative protein abundance, protein phosphorylation, and **(C)** mRNA data of A549 wt cells synchronously stimulated with 220 ng 5’ppp-dsRNA were used to set up and calibrate a dynamic mathematical model. Dots represent quantitative data of four **(B)** or two **(C)** biologically independent experiments, lines represent average model fit and dashed lines represent the confidence interval. **(D)** IFNB1 mRNA expression kinetics upon dsRNA electro-transfection was analyzed in A549 wt, A549 IRF3 KO, and A549 ROSA26-IRF3 (A549 IRF3 KO ROSA26-IRF3) expressing cells and used for model validation. Values were normalized to the housekeeping gene GAPDH and the 0 hour time point subsequently using the 2^-ΔΔCt^ method. Graph depicts mean ± SD of three biologically independent experiments. **(E)** Core model simulation of IFNB1 mRNA expression in reduced IRF3 protein level conditions.

In order to challenge the model with conditions not used for its training, we performed additional experiments in A549 cells with artificially lowered concentrations of IRF3. For this purpose, we used A549 IRF3 KO cells and restored IRF3 to approximately 5% of wildtype level by lentiviral transduction using a weak promoter (murine ROSA26) (Supplementary Figure S2A). A549 wt, A549 IRF3 KO, and A549 ROSA26-IRF3 cell lines were synchronously stimulated by electro-transfection with 5’ppp-dsRNA and IFNB1 mRNA expression was monitored over time. As expected, IRF3 knockout diminished IFNB1 induction by more than 1000-fold (Figure 3D). A549 ROSA26-IRF3 cells showed an intermediate phenotype with slower induction kinetics and >1 log10 reduced transcript levels at four hours post stimulation (Figure 3D). Notably, this dampened induction kinetics was correctly captured by our model after adjusting only the IRF3 concentration, accurately predicting the qualitative changes, with even the quantitative predictions matching the actual measurements reasonably well (Figure 3E). For further model validation, we simulated the fraction of nuclear IRF3 upon stimulation with varying concentrations of 5’ppp-dsRNA (Supplementary Figure S3), compared these model predictions to experimental data (Figure 1C), and observed a high agreement between experimentally obtained data and model simulation. This validation using independent data highlighted the usability of our ODE model not only to meaningfully fit measured data, but also to profoundly predict the outcome of signaling under perturbed conditions and, hence, to draw functional conclusions from *in silico* simulations.

### IFN feedback upon RLR stimulation is essential for high and sustained ISG expression, but not for IFN production

IFN-exposure and signaling downstream its cognate receptor through the JAK/STAT cascade does *not* induce the production of IFN in an auto-feedback manner (Supplementary Figure S4). It does, however, upregulate numerous components required for PRR signaling, in particular RIG-I itself, thereby sensitizing cells for IFN production (indirect positive feedback). Here, we employed A549 cells with a double knockout of the type I (IFNAR1) and type III (IFNLR) IFN-receptors, termed IFN receptor double knockout (A549 IFNR DKO), to further characterize RLR signaling dynamics in the presence or absence of type I and III IFN feedback. Lacking both IFN receptors renders those cells insensitive to type I or III IFN stimulation (Supplementary Figure S2C). A549 wt and A549 IFNR DKO cells were synchronously stimulated by electro-transfection with 5’ppp-dsRNA, and type I and type III IFN induction (Figure 4A) as well as ISG mRNA expression (Figure 4B) was monitored over time. Interestingly, the dynamics of IFN mRNA induction upon stimulation was indistinguishable between both cell lines: after a rapid induction within the first hour of dsRNA stimulation (as seen in Figure 2) and reaching peak levels between six and eight hours, IFN mRNA expression decreased again between 12 and 24 hours post stimulation. This further corroborated the notion that IFN production is dependent on RLR signaling and completely independent on IFN signaling (Figure 4A). Similarly, using a multiplex immunoassay (U PLEX IFN Combo, Meso Scale Diagnostics) we analyzed IFN-β and IFN-λ1 secretion, which could be detected from six hours post electro-transfection onwards and plateaued at later time points with no notable difference between wt and IFNR DKO cells (Figure 4C). In contrast, comparing A549 wt and IFNR DKO cells, a substantial difference was observed for mRNA expression of the ISGs IFIT1 and MX1 (Figure 4B). While initial induction up to two hours (MX1) or six hours (IFIT1) was still unaffected, the lack of IFN feedback in IFNR DKO cells led to a significant decrease in IFIT1 mRNA levels over the following hours and a substantially lower overall induction for MX1 (Figure 4B). Moreover, also at the protein level MX1 was not detected in dsRNA stimulated A549 IFNR DKO cells, whereas wt cells showed increasing expression from 12 hours on (Figure 4D). In line with the IFN-dependent production of MX1, phosphorylated STAT2, a hallmark of active IFN signaling and ISGF3 formation, was undetectable in A549 IFNR DKO cells. Interestingly, key components of RLR signaling, such as RIG-I or CCL5 expression, IRF3 phosphorylation, and IkBα degradation, showed virtually identical kinetics in wt and IFNR DKO cells (Figure 4D). This clearly corroborates that IFN production is solely dependent on RLR signaling and not (directly) dependent on feedback via IFN receptor signaling. As RIG-I expression was considerably upregulated over time even in IFNR DKO cells, this furthermore demonstrates that the RLR pathway exhibits an IFN-independent feedback regulation through which the pathway reinforces itself, e.g., by upregulation of the sensor RIG-I.

**Figure 4.**
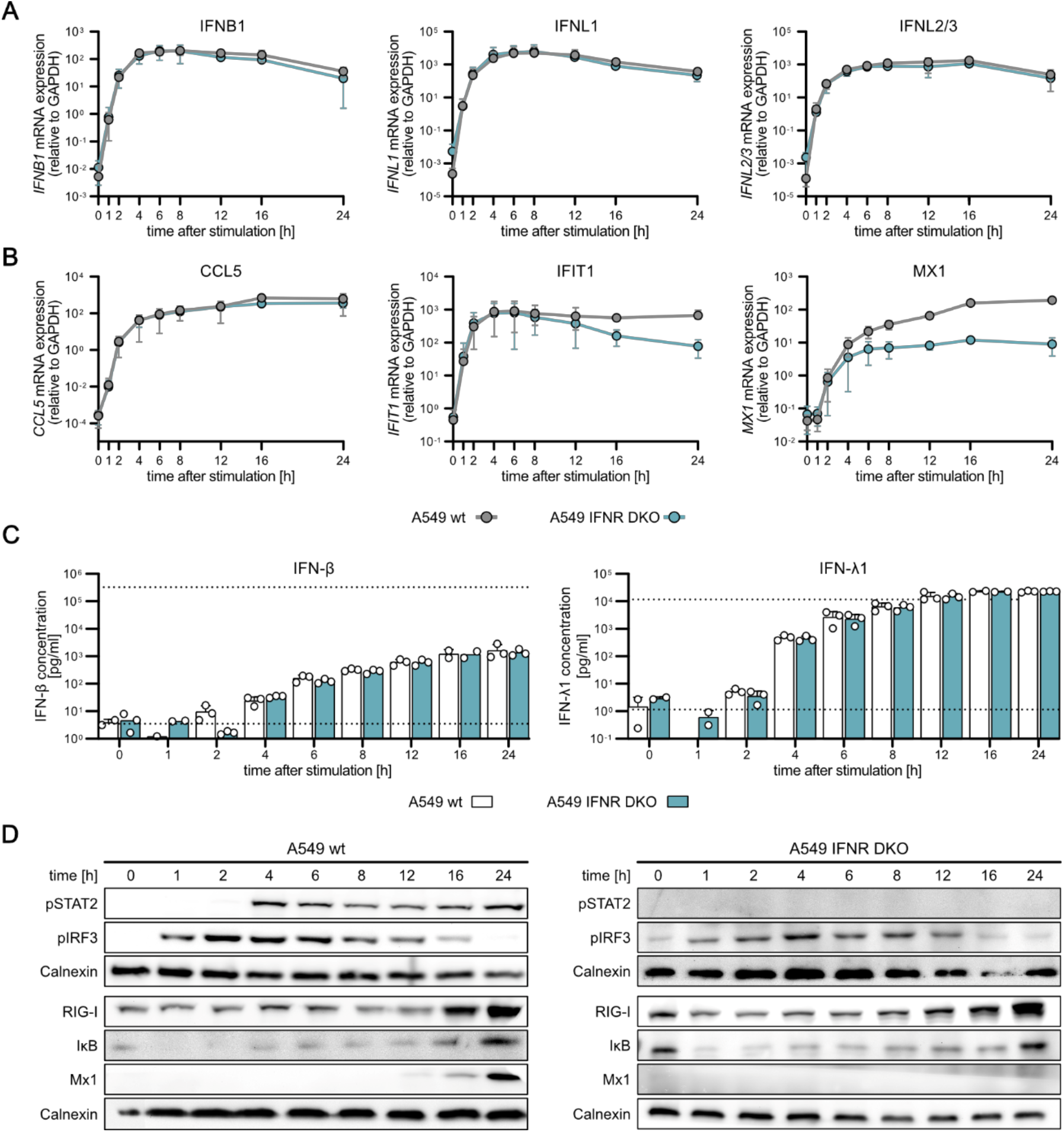
RIG-I signaling kinetics impact on IFN signaling. A549 wt and an A549 IFNAR1 IFNLR double knockout cell line (IFNR DKO) were synchronously stimulated by electro-transfection with 220 ng in vitro generated 5’ppp-dsRNA. **(A)** IFNB1, IFNL1 (IL-29), and IFNL2/3 (IL-28) or **(B)** CCL5, IFIT1, and MX1 mRNA expression kinetics was analyzed using quantitative RT-PCR. Values were normalized to the housekeeping gene GAPDH by applying the 2^-ΔCt^ method. **(C)** Secreted IFN-β and IFN-λ1 protein concentrations in pg/ml were determined using a multiplex immunoassay (U-PLEX IFN Combo, Meso Scale Diagnostics) in A549 wt and A549 IFNR DKO cells. Dashed lines indicate upper and lower limit of quantification, respectively. **(D)** Protein abundance and phosphorylation status kinetics of key signaling components in A549 wt and A549 IFNR DKO cells using western blot analysis. Graphs depict mean ± SD of three biologically independent experiments (A-C) or representative blots of two biologically independent experiments (D).

### Expanded RIG-I pathway model comprising IFN feedback accurately reproduces the full cell-intrinsic antiviral response

Having established a dynamic pathway model of the core RIG-I signaling module, we were able to accurately predict the pathway outcome from viral RNA recognition by RIG-I to transcription of IFN-β by IRF3 within the first few hours after RIG-I stimulation (Figure 3). However, so far, our model did not capture the translation and secretion of IFNs and, thus, could not recapitulate the effects of IFN signaling that considerably shape the antiviral state over the prolonged time of RLR stimulation (e.g., 24 hours, Figure 4). In order to overcome this limitation, we extended our model by combining our core RIG-I signaling module with an ODE-based model of the type I IFN-triggered JAK/STAT signaling pathway previously developed by Maiwald and colleagues [24]. The latter is a detailed model of JAK/STAT signaling upon stimulation with IFN-α. To connect the two modules, we quantified secreted IFN levels upon RIG-I pathway stimulation and linked its production to the IFN-β mRNA levels. The produced IFN levels acted as the input for the IFN signaling module from literature (Figure 5A, B). Our combined model comprises the full cell-intrinsic antiviral response from incoming viral RNA to the expression of antiviral effector proteins downstream of IFN signaling. The JAK/STAT model introduced 66 additional species, engaged in 41 reactions. Importantly, we fixed all rate constants except for one to the values determined previously– for the RIG-I module as described above, and for the IFN signaling module as determined by Maiwald et al. [24]. The single adjusted rate constant k_68_ links the amount of produced IFN to the effective concentration triggering IFNAR signaling. Protein concentrations of components in the JAK/STAT pathway were further updated to values obtained in A549 cells if available to account for cell specific differences in protein synthesis. Without adjusting any further parameter, the model was able to accurately describe all measured pathway components over the course of the experimental time frame of 24 hours (Figure 5C). One component of the model that suffered high experimental variability was phosphorylation of STAT2 due to its high sensitivity to even very low concentrations of IFN, resulting in a very pronounced initial activation. To further account for a more detailed description of ISG expression, the model was extended to include the dynamics of selected ISG mRNAs and proteins. Eight new species were introduced and the differential equation for RIG-I was updated to account for its upregulation (Supplementary Material).

**Figure 5.**
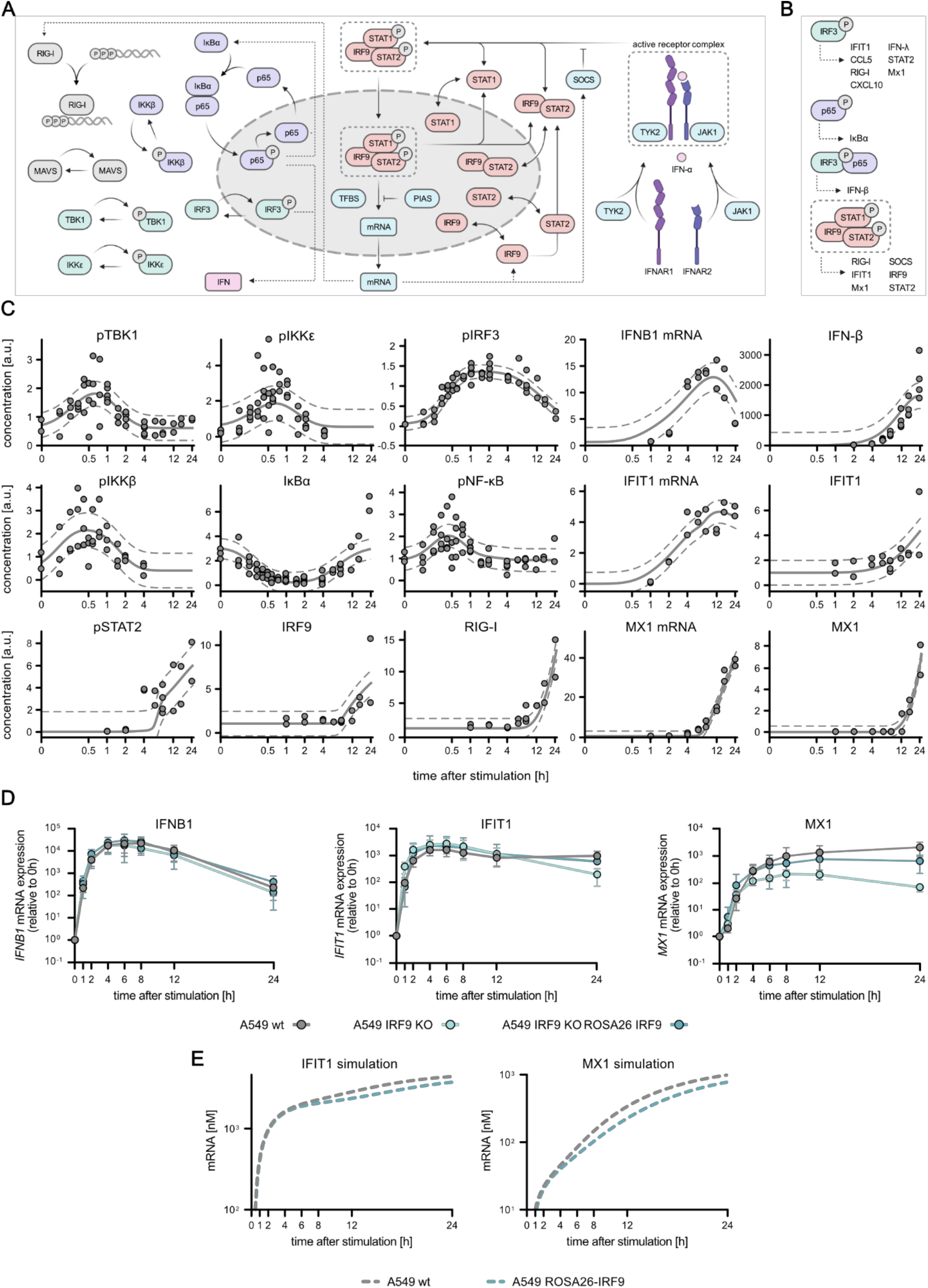
Dynamic pathway model of core RIG-I signaling coupled to IFN signaling model. Extension of the RIG-I signaling pathway mathematical model to include IFN signaling by using quantitative mRNA expression, protein phosphorylation, and protein abundance data obtained upon synchronous stimulation with 220 ng *in vitro* generated 5’ppp-dsRNA. **(A)** Schematic depiction of the established core RIG-I signaling model coupled to a previously published mathematical model of IFN signaling [24]. **(B)** Gene expression legend for different transcription factors used in the extended mathematical model. **(C)** Quantitative protein abundance, protein phosphorylation, and mRNA data were used to combine and calibrate the extended, dynamic model of the full antiviral system. Dots represent quantitative data of two biologically independent experiments, lines represent average model fit and dashed lines represent the confidence interval. **(D)** IFNB1, IFIT1, and MX1 mRNA expression kinetics upon synchronous dsRNA stimulation was analyzed in A549 wt, A549 IRF9 KO, and A549 ROSA26 IRF9 (A549 IRF9 KO ROSA26-IRF9) expressing cells and used for model validation. Values were normalized to the housekeeping gene GAPDH and the 0 hour time point subsequently using 2^-ΔΔCt^. Graphs depict mean ± SD of three biologically independent experiments. **(E)** Coupled model simulation of IFIT1 and MX1 mRNA upon varying IRF9 protein levels.

The downstream production of antiviral effector proteins, such as MX1, could be reproducibly quantified experimentally and very accurately captured by the model (Figure 5C). In their study, Maiwald and colleagues identified IRF9 as a central and rate-limiting component of IFN signaling [24]. Hence, in order to further challenge and validate our model, we experimentally modulated IRF9 levels. As in our A549 cell system IRF9 appeared not to be limiting (Supplementary Figure S5), we decreased its expression by CRISPR/Cas9-mediated KO (IRF9 KO) and subsequently expressed IRF9 from the weak murine ROSA26 promoter (ROSA26-IRF9) using lentiviral transduction (Supplementary Figure S2E). We synchronously stimulated cells by electro-transfection of 5’ppp-dsRNA and measured mRNA expression of IFNB1, IFIT1, and MX1 over time (Figure 5D). Very similarly to IFNR DKO conditions (Figure 4), IFNB1 mRNA levels were exclusively controlled by RLR downstream signaling and, hence, not sensitive to modulation of IRF9 levels, whereas IFIT1 was slightly affected and the “classical” ISG MX1 was considerably impacted by lacking or reduced IRF9 (Figure 5D). In essence, these results indicate that IRF9 levels are critical for the expression of some genes (IFIT1, MX1), whereas others seem to be completely independent of IRF9 protein levels (IFNB1). The measured outcome of this experimental perturbation of the signaling system was correctly predicted by our model, further corroborating its robustness and reliability (Figure 5E).

### RIG-I signaling dynamics can be accurately modelled in different cell lines

For model establishment we focused on A549 cells for their well-known capacity for RLR and IFN signaling, and their amenability to genetic modification. As innate immune signaling pathways are evolutionary ancient and well-conserved, we tested our model’s potential to be adapted to other cell types. For this purpose, we performed side-by-side 5’ppp-dsRNA stimulation of A549 cells and an unrelated liver (hepatocellular carcinoma) cell line, HepG2, and compared the ensuing mRNA expression (Figure 6A, B), protein abundance and phosphorylation (Figure 6C), as well as IFN secretion (Figure 6D) dynamics. Surprisingly, whereas IFIT1 and CCL5 mRNA expression resembled similar kinetics in both cell lines, IFN mRNA expression (Figure 6A) as well as IFN secretion (Figure 6D) was significantly reduced in HepG2. Accordingly, IFN receptor signaling-dependent STAT2 phosphorylation and subsequent RIG-I and MX1 mRNA and protein expression were strongly reduced in HepG2 (Figure 6B, C). Notably, although IFIT1 mRNA expression kinetics (Figure 6A) was identical in both cell lines, protein expression was clearly reduced in HepG2 cells as compared to A549 (Figure 6C). We next adapted our mathematical model to the new cell line, considering the differential protein expression levels in this cell line. In order to do so, we measured the base-line protein concentrations of the major signaling components in HepG2 cells by mass-spectrometry and label-free quantification and adjusted the respective initial concentrations in our model (Supplementary Material). Strikingly, adapting these initial conditions and without any further adjustment of kinetic rate constants, the model was capable to adequately describe the mRNA kinetics of IFNB1 and MX1 in HepG2 cells (Figure 6E). Only experimental results of IFIT1 mRNA kinetics did not fully resemble the reduced effect in model simulations (Figure 6E). This highlights the potential of our model to be adapted to different cell lines by only adjusting the base-line expression levels of signaling components and corroborates the notion that RLR and IFN signaling are highly conserved and robust systems.

**Figure 6.**
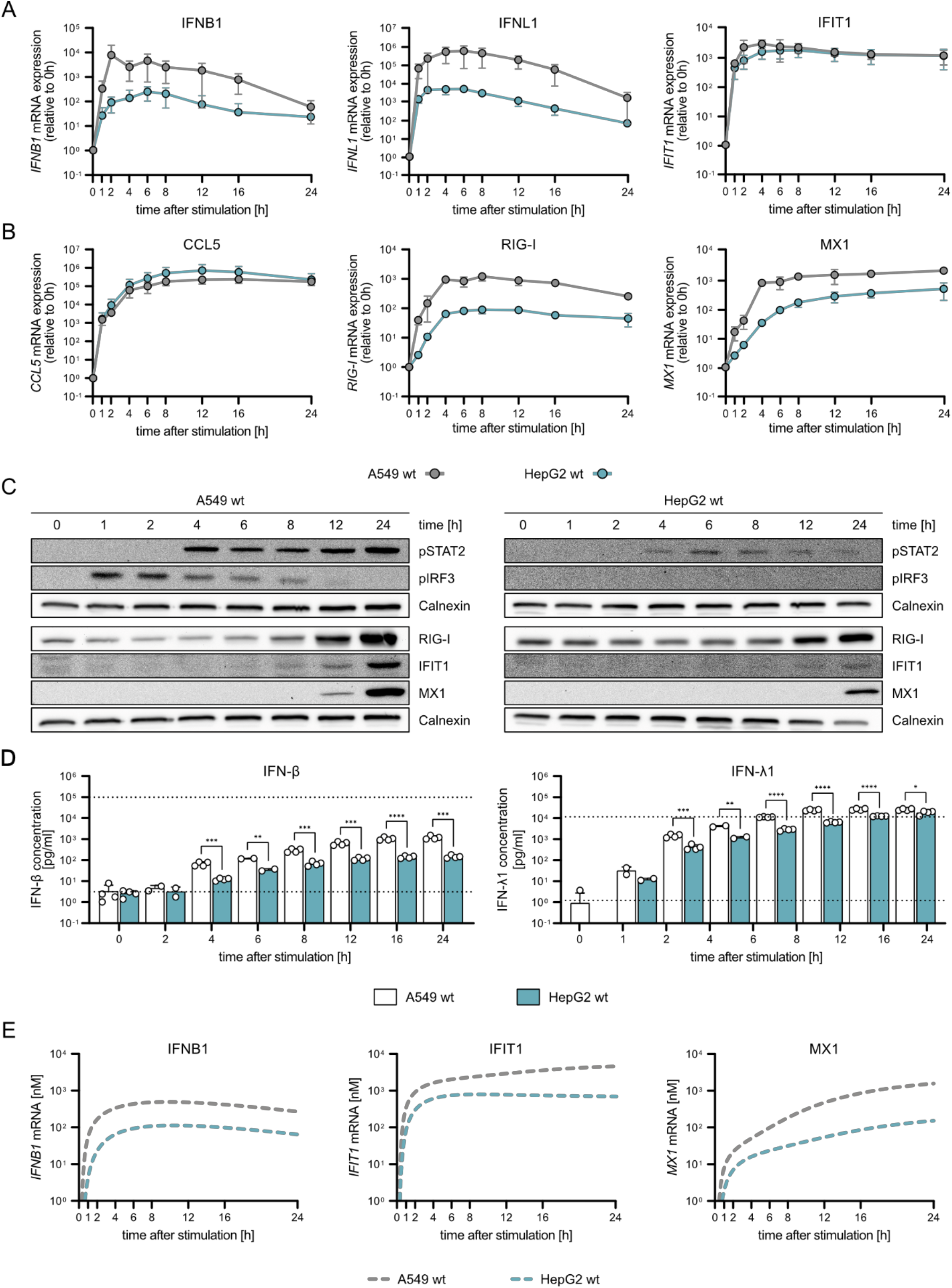
Characterization of RIG-I signaling dynamics upon synchronous dsRNA simulation in two unrelated cell lines. A549 wt and HepG2 wt cells were synchronously stimulated with 220 ng 5’ppp-dsRNA by electro-transfection and RNA, protein, and supernatants were harvested at different time points. **(A)** IFNB1, IFNL1, IFIT1 and **(B)** CCL5, RIG-I, MX1 mRNA expression kinetics in A549 wt and HepG2 wt cells was analyzed using quantitative RT-PCR. Values were normalized to the housekeeping gene GAPDH and the 0 hour time point subsequently using 2^-ΔΔCt^. **(C)** Quantitative protein abundance and protein phosphorylation upon synchronous stimulation in A549 wt and HepG2 wt cells was analyzed using western blot analysis. **(D)** Secreted IFN-β and IFN-λ1 protein concentrations were determined using a multiplex immunoassay (U-PLEX IFN Combo, Meso Scale Diagnostics) in A549 wt and HepG2 wt cells. Dashed lines indicate upper and lower limit of quantification, respectively. **(E)** Model predictions for IFNB1, IFIT1, and MX1 mRNA expression upon synchronous stimulation of A549 wt and HepG2 wt cells. Graphs depict mean ± SD (A, B, D) or representative blots (C) of three biologically independent experiments. Statistical significance was determined with Student’s t-test (****: p < 0.0001, ***: p < 0.001, **: p < 0.01, *: p < 0.05).

### The impact of viral antagonists on antiviral signaling dynamics can be properly simulated by the established mathematical model

Upon virus infection, kinetics and magnitude of the ensuing IFN response critically determine the outcome of infection. Therefore, most viruses have evolved potent antagonists capable of inhibiting host-cellular antiviral responses. Whereas most studies focus on the degree of IFN inhibition at a fixed (late) time-point in the context of strong and constitutive overexpression of the viral antagonist, in a natural infection, viral proteins get enriched over time and may lead to a delay of the IFN response. To better understand these critical host-virus interactions it thus is important to investigate the effect of viral immune antagonism at a dynamic level. Our model of the IFN system may offer a valuable tool for studying the impact of viral antagonists onto the dynamic profile of ISG induction. In order to test this, we have chosen a couple of well-described viral proteins with different modes of action: NS3/4A of hepatitis C virus (HCV) proteolytically inactivates the central adapter MAVS [15], N^pro^ of classical swine fever virus (CSFV) triggers the degradation of IRF3 [13,14], and NS5 of dengue virus (DENV) is described to degrade STAT2 [32,33]. While NS3/4A and N^pro^ target RLR signaling and thereby IFN induction, NS5 rather blocks signaling downstream of the IFN receptor.

Using lentiviral transduction, we generated A549 cells stably expressing varying levels of the viral proteins. In case of NS3/4A, already minute amounts sufficed for maximum MAVS cleavage. Therefore, we here used graded doses of Simeprevir, a pharmacological compound specifically inhibiting NS3/4A protease activity [34,35], which allowed us to mimic the effect of decreasing NS3/4A activity. Cells were then stimulated by electro-transfection of 5’ppp-dsRNA and IFIT1 mRNA levels were measured over the course of 24 hours (Figure 7). For NS3/4A we observed a very efficient cleavage of MAVS (Figure 7B), leading to a substantial (30-fold) reduction of IFIT1 mRNA levels particularly at early time points (Figure 7A). Notably, a basal level of MAVS withstood NS3/4A expression (Figure 7B), leading to remaining signaling and induction of IFIT1 expression. As expected, increasing concentrations of Simeprevir dose-dependently decreased NS3/4A activity leading to increased amounts of intact MAVS protein (Figure 7B). This dose-dependent effect was reflected in IFIT1 kinetics, with increasing concentrations of Simeprevir restoring IFIT1 induction to the level of the catalytically inactive NS3/4A S139A control (Figure 7A). Taken together, increasing NS3/4A activity strongly reduced expression of IFIT1. This effect was less pronounced for later time points (12 and 24 hours), nonetheless, the overall kinetics of induction was not altered significantly. Interestingly, this expression dynamics was also qualitatively predicted by our mathematical model when we systematically reduced MAVS concentrations *in silico* (Figure 7C), highlighting the utility of the modeling approach to investigate viral immune evasion.

**Figure 7.**
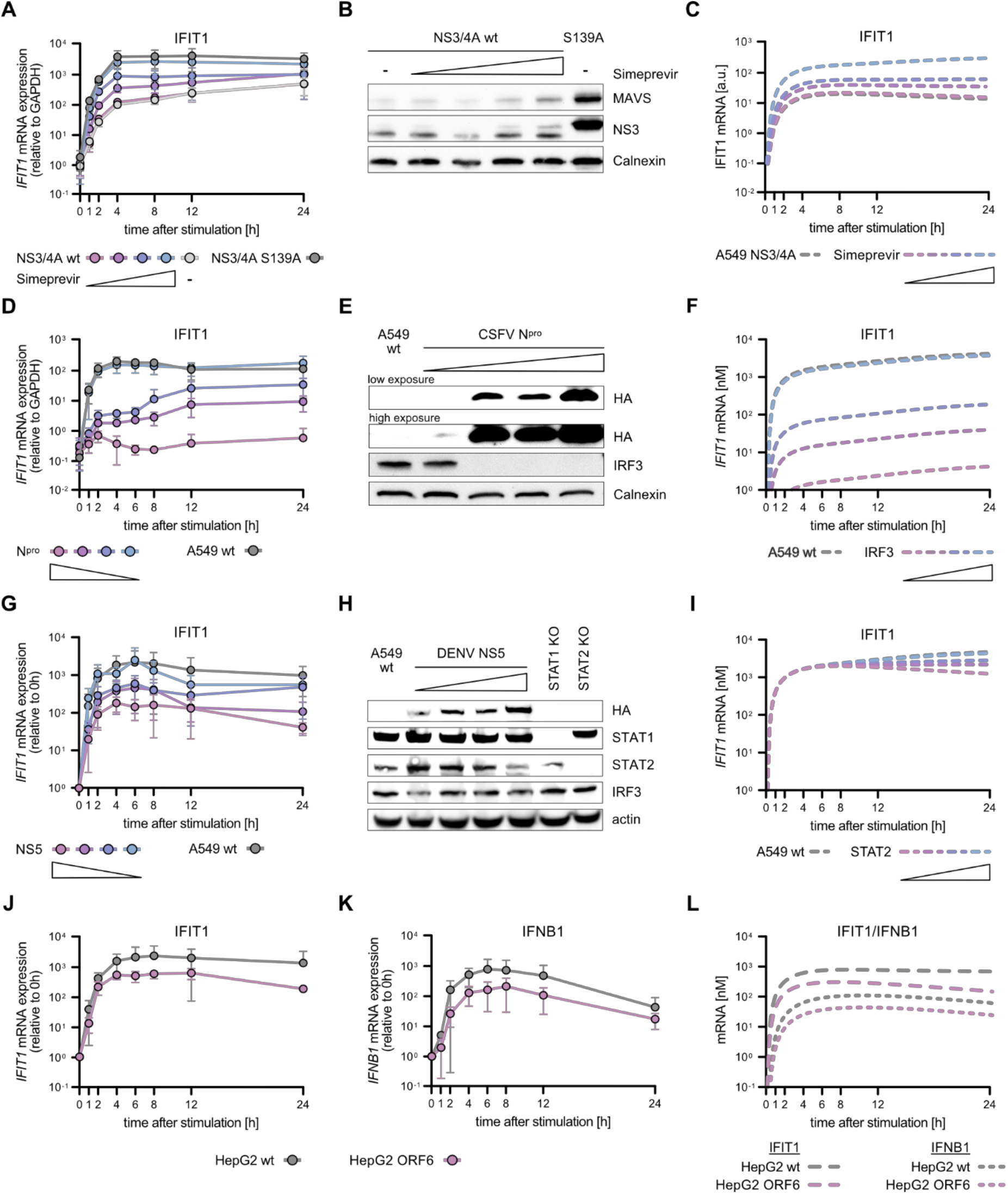
Influence of various viral antagonists on RIG-I signaling dynamics in A549 and HepG2 cells. A549 cells stably expressing the viral proteins NS3/4A of hepatitis C virus (HCV, A-C), N^pro^ of classical swine fever virus (CSFV, D-F), NS5 of dengue virus (DENV, G-I) or HepG2 cells stably expressing ORF6 of SARS coronavirus type 2 (SARS-CoV-2, J-L) were generated using lentiviral transduction. **(A-C)** A549 cells stably expressing either the HCV NS3/4A protein or a catalytically inactive version (NS3/4A S139A) were treated with increasing protease inhibitor (Simeprevir) concentrations overnight and subsequently synchronously stimulated with 220 ng 5’ppp-dsRNA. **(A)** IFIT1 mRNA expression kinetics and **(B)** basal NS3 and MAVS protein levels were analyzed in A549 NS3/4A wt and A549 NS3/4A S139A overexpression cell lines. **(C)** Model simulation of IFIT1 mRNA kinetics upon synchronous stimulation with dsRNA in A549 cells expressing NS3/4A in the absence or presence of increasing amounts of Simeprevir. **(D-F)** A549 wt cells and cells expressing different levels of HA-tagged CSFV N^pro^ were synchronously stimulated with 220 ng 5’ppp-dsRNA. **(D)** IFIT1 mRNA expression kinetics and **(E)** basal HA-N^pro^ or IRF3 protein levels were analyzed in A549 wt or N^pro^ overexpression cell lines using qRT-PCR and western blotting. **(F)** Model simulation of IFIT1 mRNA kinetics upon synchronous stimulation with dsRNA with varying levels of IRF3. **(G-I)** A549 wt cells and cells expressing different levels of HA-tagged DENV NS5 were synchronously stimulated with 220 ng 5’ppp-dsRNA. **(G)** IFIT1 mRNA expression kinetics and **(H)** basal HA-NS5, STAT1, STAT2, and IRF3 protein levels were analyzed in A549 wt, STAT1 KO, STAT2 KO or NS5 overexpression cell lines. **(I)** Model simulation of IFIT1 mRNA kinetics upon synchronous stimulation with dsRNA with varying levels of STAT2. **(J)** IFIT1 and **(K)** IFNB1 mRNA expression kinetics were analyzed in HepG2 wt cells and cells expressing Strep-tagged SARS-CoV-2 ORF6 upon synchronous stimulation with 220 ng 5’ppp-dsRNA. **(L)** Model simulation of IFIT1 and IFNB1 mRNA expression upon synchronous stimulation. qRT-PCR values were either only normalized to the housekeeping gene GAPDH (A, D) or additionally normalized to the 0 hour time point (G, J-K). Graphs depict mean ± SD (A, D, G, J-K) or representative blots (B, E, H) of three biologically independent experiments.

N^pro^ directs the important transcription factor IRF3 for degradation (Figure 7E), and accordingly we observed a near complete inhibition of IFIT1 induction for the highest N^pro^ levels (Figure 7D). With decreasing expression of the viral antagonist, IFIT1 induction dose-dependently recovered (Figure 7D, E). Very interestingly, although similar to NS3/4A it targets the RLR cascade at a step upstream of IFN expression, in contrast to the HCV protease, N^pro^ considerably affected the kinetic profile of IFIT1 induction, very strongly repressing the early expression up to eight hours post stimulation (Figure 7D). We then used our model to simulate the N^pro^ effect by gradually reducing IRF3 abundance in our model. The predicted kinetics of the antiviral response matched our experimental data at early time points (Figure 7F, compare to Figure 7C) and our results underscore the potency of N^pro^ as an immune antagonist. Next, we tested the effects of DENV NS5 expression, which targets STAT2 for degradation and therefore affects signaling downstream of the IFN receptor, but not IFN induction through the RLR pathway. Increasing levels of NS5 indeed led to decreasing amounts of STAT2, but not STAT1 or IRF3 (Figure 7H, Supplementary Figure S2B). Of note, even for the highest concentration of NS5 STAT2 degradation was only partial (approx. 40 % remaining). Correspondingly, IFIT1 expression was affected particularly at late time points when its transcription is predominantly driven by IFN signaling, leading to a decrease in mRNA levels from eight hours post stimulation onwards (Figure 7G). Despite appreciable amounts of remaining STAT2, the degree of this decrease was very comparable to the condition of IFNR DKO (Figure 4B) or IRF9 KO (Figure 5D). Surprisingly, at least for the higher concentrations of NS5, also early induction of IFIT1 was clearly inhibited, even to an extent comparable to NS3/4A (Figure 7A). This was also not predicted when we simulated IFIT1 induction under conditions of limited STAT2 availability in our model (Figure 7I). These findings strongly suggest NS5 may have additional functions in the early stages of the antiviral response that have not been described before.

Finally, we used a viral antagonist with more than one defined target: ORF6 of the recently emerged SARS coronavirus 2 (SARS-CoV-2). ORF6 was shown to negatively impact both IFN induction and IFN signaling [36], likely by blocking IRF3 [37,38], STAT1, and STAT2 [39] nuclear translocation. Surprisingly, in A549 cells ORF6 expression did not impact IFIT1 or IFNB1 expression upon dsRNA stimulation (Supplementary Figure S6, Supplementary Figure S2D). While we cannot explain this effect in A549 cells, we could observe a clear effect in HepG2 cells (Figure 7J, K, Supplementary Figure S2D). In HepG2 cells, ORF6 affected IFIT1 mRNA induction qualitatively similar to NS5, with a moderate reduction at early time points, but a clear decrease between eight and 24 hours post stimulation. These findings are consistent with model simulations under conditions of simultaneously reduced IRF3 and STAT2 levels (Figure 7L).

In summary, we established an ODE-based mathematical model of antiviral signaling, comprising the RLR pathway of IFN induction, as well as IFN signaling downstream of the IFN receptor. This model can very accurately simulate the activation of individual signaling components over time, as well as the induction of ISGs and the production of antiviral effector proteins. Importantly, it permits *in silico* simulation of viral interference with this cell-intrinsic defense system and offers a valuable tool to study the impact of yet unknown viral antagonists or other factors perturbing RLR and/or IFN signaling.

## DISCUSSION

All nucleated cells possess defense systems to protect themselves against infection by microbes such as viruses. While higher organisms have furthermore evolved sophisticated adaptive immune systems, evolutionary ancient innate immune responses [40,41] even in humans still represent the vital basis for all higher tiers of immune reactions. For one, cytokines produced and secreted at early stages of infection play a critical role in proper launching of adaptive immune responses and their coordination [42]. But even beyond activating and coordinating the action of professional immune cells, immediate cell-intrinsic responses of the infected cells themselves potently suppress microbial, in particular viral replication, which frequently is an absolute prerequisite for successful clearance of the pathogen [43]. Very recently, it has been shown for SARS-CoV-2 that improper functioning of the IFN system very strongly predisposes individuals for life-threatening COVID-19 [44–46], and we and others have shown that a particularly swift and strong IFN response in children associates with their striking resilience towards severe courses of the disease [47,48]. In general, the immune-driven pathogenesis of COVID-19 impressively illustrates the critical importance of timely and well-coordinated cytokine production for a successful immune response, and the potential of a dysregulated IFN and cytokine system to cause substantial pathology [49]. Lastly, slowed-down or dampened IFN responses are observed in elderly individuals as part of the so-called immunosenescence, rendering aged patients more vulnerable to infectious diseases [50,51], but also occur at conditions of lower ambient temperature, likely contributing to the seasonality of certain viral infections [52–54].

The emerging importance of pathway dynamics and the difficulty in simultaneously activating the RIG-I pathway prompted us to optimize stimulation conditions and characterize the RIG-I signaling kinetics more precisely. In fact, we observed a striking difference in the activation kinetics of RIG-I signaling when we used electroporation-based millisecond-transfection as opposed to classical liposome- and endocytosis-based transfection. We observed a virtually synchronous activation of all cells as determined by IRF3 nuclear translocation, occurring already 15 to 30 minutes after stimulation. The first transcripts of type I and III IFNs could be detected after 45 to 60 minutes, with measurable amounts of secreted IFNs in the culture supernatants after four to six hours. In order to further substantiate our kinetic understanding of the pathway, we used mathematical modelling. We used literature knowledge of the core pathway topology to set up a set of ODEs mechanistically representing all major canonical signaling steps. We have previously modelled dsRNA recognition by RIG-I and its downstream signaling, however, with a focus on the quantitative output rather than signaling kinetics [55]. Another study also modelled RIG-I (and TLR3) signaling [28], even using electro-transfection of dsRNA similar to our approach, however, with a focus on cross-talk to TLR signaling via TRAF molecules and considerably less temporal resolution in the experimental data. A central feature of the antiviral system is strong positive feedback through IFN. Albeit RLR signaling itself is known to induce certain antiviral effectors (ISGs) directly, such as IFIT1 [30,56,57], the full-fledged transcriptional program mediating the antiviral state of a cell is only established downstream of IFN receptor signaling. Importantly, also the RLRs themselves– RIG-I, MDA5, and LGP2– as well as the IRF3-like transcription factor IRF7, are ISGs substantially induced upon IFN signaling. In an actual viral infection, autocrine IFN signaling, hence, reinforces viral sensing and IFN production. In our experimental system, this increased RNA sensing and IFN production plays less of a role, as the dsRNA is only pulse-transfected and not replicated for prolonged periods of time as would be the case in an infection. In fact, when we used “IFN-blind” cells, harboring a functional double knockout of the type I and type III IFN receptors (IFNR DKO), the induction dynamics of IFNs was not affected at any time point tested. In contrast, IFIT1, being induced by both IRF3 and IFN signaling [30,56,57], was affected only at later time points and MX1, canonically strictly dependent on IFN signaling [1,58–60], was substantially impacted throughout. Nevertheless, MX1 was still significantly induced even in the absence of IFN signaling, which has not been broadly appreciated previously, but in fact has been shown in human fibroblasts upon HCMV infection [61]. The central and essential feedback of the RLR system through IFN must be taken into account when considering the full cell-intrinsic antiviral response, in particular for timeframes of more than four hours (after which secreted IFN can be detected). We, therefore, coupled our mathematical model to a previously published model of type I IFN signaling [24]. To couple both models, we took the IFN produced as the major output of our core RIG-I module and used it as the input for the IFN signaling module, only applying one fitted scaling factor. The IFN model is also based on ODEs and was trained on quantitative immunoblotting data, similarly to our RIG-I model. However, experimentally that study used IFN-α for stimulation, which is (at least initially) not produced downstream of RLRs in our cell system. Nonetheless, it is a type I IFN much like IFN-β and potentially does not affect signaling dynamics to a relevant extent. A further difference to our system is the employed cell line, which was the human hepatoma cell line Huh7.5 in that study. Reassuringly, despite the two models were fitted to data from different cell lines, all kinetic rate constants of the published IFN model could be employed in our combined model of the full antiviral pathway. Upon only adjusting the initial protein concentrations to the ones measured in our A549 cells, the combined model of RIG-I and IFN signaling with these parameters very accurately described our measured 24 hour time course data. This impressively highlights the substantial degree of conservation of this pathway across different cell types, which we have also previously observed between two different hepatoma cell lines and primary human hepatocytes [62]. To further corroborate the cell line independence of our comprehensive model of antiviral signaling, we validated it using time course measurements in the unrelated HepG2 cell line. Again, upon only adjusting the cell line-specific initial protein concentrations, our combined model was able to adequately predict RLR as well as IFN signaling dynamics in HepG2 cells upon electro-transfection of 5’ppp-dsRNA. We, hence, expect that our model is generalizable to a wide variety of different cell lines and types by simply adjusting the basal protein concentrations of the proteins involved in signaling. This advertises the model as a valuable tool to study kinetic aspects of antiviral signaling even with experimental measurement only at a limited number of time points. We cannot rule out, however, that in some cell types– e.g., myeloid cells of the professional immune system– further regulatory systems not represented in the model might be in place, or additional transcription factors of (partially) overlapping function may exist that require modifications to the model topology. One such example is IRF7, a transcription factor highly similar to IRF3, which is expressed to high levels particularly in plasmacytoid dendritic cells, rendering them competent to produce excessive amounts of IFN, including INF-α, even prior to the positive feedback through the JAK/STAT pathway [63–65].

We propose that our dynamic pathway model additionally offers a powerful framework to study virus-host interactions. As a proof of principle, we have selected four well-known viral proteins interfering at defined steps with the host-cell antiviral defense: the proteases of hepatitis C virus (NS3/4A) and CSFV (N^pro^) both target RLR signaling and, thus, inhibit the induction of IFN, whereas dengue virus NS5 interferes with IFN signaling. Lastly, we picked the ORF6 protein of the novel coronavirus SARS-CoV-2, which exhibits a multi-pronged strategy to inhibit the antiviral system both at the induction as well as the effector level. NS3/4A is a vital protease involved in maturation of the HCV non-structural proteins, but it also very efficiently cleaves and inactivates MAVS [15]. We found its activity to have an overall dampening effect on IFIT1 induction, most pronounced for the very early time points, coherent with its targeting the very early induction phase of the antiviral response. CSFV N^pro^ directs IRF3 for proteasomal degradation [13,14], thus also targeting early IFN production. In contrast to NS3/4A, though, it leaves NF-κB signaling unhindered, and should, hence, allow for the production of certain pro-inflammatory cytokines [66–68]. In our experiments, we found a substantially stronger impact of N^pro^ on early IFIT1 induction, with a particularly pronounced effect setting in after two hours. When we simulate the effects of these two viral antagonists in the model, it very closely captures the impact of both NS3/4A and N^pro^. Complementary to NS3/4A and N^pro^, dengue virus NS5 targets signaling downstream of the IFN receptors and leads to the degradation of STAT2 [32,33]. Accordingly, we observed less of an effect for the early time points, but a clear tendency for IFIT1 levels to decrease between eight- and 24 hours, when its transcription is largely shifted from the early IRF3 to the IFN-dependent transcription factor ISGF3. Nonetheless, NS5 expression had a clear effect also at time points before eight hours, which could also not be explained by simulating a pure STAT2-mediated effect in our model. This suggests that DENV NS5 may have additional targets, potentially already in the induction phase of the antiviral defense, at or downstream of the RLRs. In fact, one previous study proposed an interaction of NS5 and the Daxx protein [69]. Here, NS5 is competing with the NF-κB/Daxx interaction, hence, leading to NF-κB release and the subsequent expression of RANTES [69,70]. It will be interesting to follow up on this in future studies. Lastly, we expressed SARS-CoV-2 ORF6. Here, we had to resort to HepG2 cells, as in A549, for reasons unknown to us, ORF6 had no impact on antiviral signaling despite being reported to be amongst the strongest SARS-CoV-2 antagonists in 293T cells [36]; this phenomenon may warrant further investigation. In HepG2, ORF6 had a very modest impact at time points prior four hours post dsRNA stimulation, but subsequently limited the peak levels of IFIT1 mRNA and even led to decreasing transcript levels between eight and 24 hours. This confirms that ORF6 has several targets in the antiviral system [36], but indicates the dominant effect may be on signaling events downstream of the IFN receptor(s). This was corroborated by modelling, where inhibition of ISGF3 formation approximated the experimental data much closer than limiting IRF3 activity (Figure 7L, compare to Figure 7F).

As a future perspective, it will be highly exciting to combine our comprehensive model of innate immune signaling with models of viral replication in order to gain a better mechanistic understanding of these intricate virus-host interactions decisive for the development of disease: viral RNA, amplified rapidly during viral replication, is the inducer of antiviral signaling; antiviral signaling, via IFN, will potently suppress viral replication (i.e. suppress production of stimulatory RNA); viral RNA at the same time is translated to proteins inhibiting antiviral signaling. We have previously approached modelling the interaction between HCV and innate immune signaling, however at a much lower level of mechanistic detail [71]. Others have modelled pathogen-host interaction at various levels as well [27,72–75]. Employing our molecularly detailed model of antiviral signaling will enable a substantially higher degree of interpretability and applicability of such approaches in the future.

In summary, we here present a highly accurate characterization of the dynamics of cell-intrinsic innate immune signaling to virus infection. Using IFN-blind KO cells, we were able to dissect the induction phase downstream of RLRs and the effector phase downstream of IFN receptors, which normally are tightly linked and overlapping. We further used our quantitative time-series data to set up and calibrate the – to the best of our knowledge – most comprehensive mechanistically interpretable dynamic pathway model of the RLR/IFN signaling network. This model very accurately predicts the kinetics of signaling events downstream of RNA recognition by RIG-I, including the feedback and broadening of the response by secreted IFN and JAK/STAT signaling. Owing to its mechanistic detail, the model is capable of simulating viral immune antagonism and, *vice versa*, can be used to investigate the mechanism of action of novel virus-encoded inhibitors. Combining models as ours with models of virus infection and replication will open up new and powerful avenues of studying the dynamics of virus-host interactions.

## MATERIAL AND METHODS

### Cell culture and cell line generation

A549, HepG2 and HEK 293T cell lines were cultured in Dulbecco’s modified eagle medium (DMEM high glucose, Life Technologies, Carlsbad, CA, USA), supplemented with 10 % (v/v) fetal calf serum (FCS, Thermo Fisher Scientific, Waltham, MA, USA), 1x non-essential amino acids (Thermo Fisher Scientific), 100 U/ml penicillin, and 100 ng/ml streptomycin (Life Technologies) at 37 °C, 95 % humidity, and 5 % CO_2_. Lentiviral transduction was used for the generation of A549 and HepG2 overexpression (OE) cell lines as well as for A549 knockout (KO) cell line generation. The Clustered Regularly Interspaced Short Palindromic Repeats (CRISPR)/Cas9 technology was used for stable KO generation in A549. DNA oligonucleotides coding for guideRNAs (listed in Supplementary Table S1) against the respective genes were designed with e-crisp.org [76] and cloned into the expression vector LentiCRISPRv2 (Feng Zhang, Addgene #52961). Lentiviral particles were produced by transiently transfecting HEK 293T cells with a total of 15 µg DNA consisting of the plasmids pCMV-dr8.91 (coding for HIV gag-pol), pMD2.G (coding for the VSV-G glycoprotein), and a lentiviral pWPI vector (coding for protein of interest) or the expression vector LentiCRISPRv2 (coding for guideRNA) in a 3:1:3 ratio for 8 hours using calcium phosphate transfection (CalPhos Mammalian Transfection Kit, Takara Bio Europe, Saint-Germain-en-Laye, France). Two days after transfection, supernatant was harvested, sterile filtered, and used for transduction of target cells. Transduction was carried out two times for 24 hours and transduced cells were selected with appropriate antibiotics (5 μg/ml blasticidin, MP Biomedicals, Santa Ana, CA, USA; 1 μg/ml puromycin, Sigma Aldrich; or 1 mg/ml geneticin (G418), Santa Cruz, Dallas, TX, USA). Single cell clones of successfully transduced cells were isolated, and KO or OE was validated by immunoblotting and, if appropriate, functional tests. A549 IRF3 KO, IRF9 KO, STAT1 KO, and IRF3-eGFP H2B-mCherry were reported previously [77–79].

### Live-cell imaging and quantification of IRF3 nuclear translocation

A549 cells stably expressing histone H2B-mCherry as well as eGFP-tagged IRF3 were seeded at a density of 6 × 10^4^ cells per 24-well. The next day, cells were stimulated with *in vitro* transcribed and chromatographically purified 400 bp 5’ppp-dsRNA [31] or poly(C) (Sigma-Aldrich) using Lipofectamine 2000 (Invitrogen, Carlsbad, CA, USA) following the manufacturer’s protocol or an in-well electro-transfection approach (Lonza). For the electro-transfection-based transfection of adherent cells, the 4D-Nucleofector® Y Unit (Lonza), the AD2 4D-Nucleofector™ Y Kit (Lonza), as well as a homemade cytomix (120 mM KCl, 0.15 mM CaCl_2_, 10 mM KPO_4_, 25 mM HEPES, 2 mM EGTA, 5 mM MgCl_2_, pH 7.6, added directly prior usage: 2 mM ATP, 5 mM GT) were utilized. First, the provided Dipping Electrode Array was used for a mock transfection to decrease the electro-transfection intensity in order to maximize cell survival and transfection efficiency. Next, DMEM was replaced with 350 µl cytomix and the electrode was inserted into the 24-well plate strictly avoiding the formation of air bubbles. Electro-transfection was performed using the FB-166 program and cytomix was replaced with warm DMEM subsequently. IRF3-GFP nuclear translocation and H2B-mCherry colocalization was analyzed using confocal microcopy equipped with an incubation chamber (Olympus FluoView FV1000) or monitored in short time periods using a 10 x magnification in an Incucyte®S3 Live-Cell Analysis System (Sartorius, Göttingen, Germany). Image analysis was conducted with the machine learning software ilastik [80] using pixel and object classification to distinguish background from cell nuclei and to determine IRF3-GFP and H2B-mCherry colocalization, respectively. Calculations for object classification were performed applying a size range of 60 to 500 and a threshold 0.85. Here, at least 500 and up to 2500 individual cells were analyzed for each time point in each condition.

### Synchronous stimulation using electro-transfection

Synchronous stimulation of the RIG-I pathway in A549- or HepG2-based cell lines was carried out using the Gene Pulser Xcell modular electro-transfection system (Bio-Rad) and a ShockPod™ cuvette chamber (Bio-Rad). Cell suspensions containing 4 × 10^6^ cells were centrifuged at 700 xg for 5 min, resuspended in 400 µl homemade cytomix and transferred to a 0.4 cm cuvette already containing 220 ng 5’ppp-dsRNA. Electro-transfection was performed at 150 V with exponential decaying pulse for 10 ms. Transfected cell suspensions were directly transferred to pre-warmed, antibiotic-free DMEM, washed twice in DMEM, and finally resuspended in 9.6 ml DMEM. Cells were seeded on 6-well plates using 1.2 ml washed cell suspension per well and time point.

### Quantitative immunoblotting

Synchronously stimulated cells were in-well lysed with 100 µl Laemmli buffer (16.7 mM TRIS pH 6.8, 5 % glycerol, 0.5 % SDS, 1.25 % β-mercaptoethanol, 0.01 % bromophenol blue), containing Benzonase® Nuclease for 10 min at room temperature, and subsequently incubated at 95 °C for 5 min. Protein extracts were separated on 8 – 12 % polyacrylamide gels by SDS-polyacrylamide gel electrophoresis (SDS-PAGE) and transferred to PVDF membranes (Bio-Rad, Hercules, CA, USA, 0.2 µm pore size) in a wet transfer approach using the Mini Trans-Blot® cell (Bio-Rad). Membranes were blocked in PBS-T or TBS-T complemented with 5 % (w/v) bovine serum albumin (BSA, Sigma-Aldrich, A3294-50G) for up to 3 hours at room temperature. Subsequently, membranes were incubated with PBS-T or TBS-T complemented with 5 % BSA and primary antibodies specific for β-actin (Sigma-Aldrich, A5441, 1:2000), calnexin (Enzo Life science, ADI-SPA-865-F, 1:1000), HA (Sigma-Aldrich, H3663, 1:1000), IFIT1 (Abnova, H00003434-DO1, 1:1000), IκBα (Cell Signaling, 9242, 1:1000), IRF3 (Santa Cruz, sc-9082, 1:1000), IRF9 (Abcam, ab126940, 1:1000), MAVS (Enzo Life science, ALX-210-929-C100, 1:1000), MX1 (kind gift of Georg Kochs, 1:1000), NS3 (NS3B, kind gift of Darius Moradpour, 1:1000), phospho-IKKα/β (Cell Signaling, 2078, 1:1000), phospho-IKKε (Millipore, 06-1340, 1:1000), phospho-IRF3 (Cell Signaling, 4947, 1:1000), phospho-NF-κB (Cell Signaling, 3033, 1:1000), phospho-STAT2 (Cell Signaling, D3P2P, 1:1000), phospho-TBK1 (abcam, ab109272, 1:1000), RIG-I (Adipogen, AG-20B-0009, 1:1000), STAT1 (BD, 610115, 1:1000), or STAT2 (Santa Cruz, sc-514193, 1:1000) at 4 °C over-night. Further, membranes were incubated with anti-rabbit horseradish peroxidase (HRP) (Sigma-Aldrich, A6154-5X1ML, 1:20 000) or anti-mouse HRP (Sigma-Aldrich, A4416-5X1ML, 1:10 000) for 1 hour at room temperature. For detection, Amersham ECL Prime Western Blotting Detection Reagent (ThermoFisher Scientific) was applied for 1 min and luminescence was detected using the INTAS ECL ChemoCam Imager 3.2 (INTAS Science Imaging Instruments, Göttingen, Germany). Western blot bands were analyzed and quantified using Image J (1.52e).

### RNA isolation, reverse transcription, and quantitative PCR

RNA isolation (New England Biolabs), reverse transcription (High-Capacity cDNA Reverse Transcription Kit, ThermoFisher Scientific), and quantitative real-time PCR (qRT-PCR; iTaq™ Universal SYBR® Green Supermix, Bio-Rad) were performed according to the manufacturer’s protocols, and qRT-PCR was performed on a CFX96 real-time-system (Bio-Rad). Sequences of specific exon-spanning PCR primers are listed in Supplementary Table S2. Values were normalized to the housekeeping gene GAPDH using the 2^−ΔCt^ method or fold changes were calculated relative to the 0 hour time point using the 2^−ΔΔCt^ method [81].

### Multiplex immunoassay

Quantitative measurement of secreted cytokines within supernatants was performed using the multiplex Meso Scale Diagnostics (MSD) platform. Briefly, supernatants of synchronously stimulated A549 wt, A549 IFNR DKO, and HepG2 wt cells were harvested, centrifuged, and multiplex assays were conducted according to the manufacturer’s instructions. The analytes IFN-β and IFN-λ1 were measured using the human U-PLEX Interferon Combo (Meso Scale Diagnostics, 15094K-1). Plate readout was conducted using the MESO QuickPlex SQ 120 instrument (Meso Scale Diagnostics) and measurements were evaluated using the MSD Discovery Workbench software.

### Protein copy number estimation of A549 and HepG2 cells by total proteome analysis using LC-MS/MS

A549 cells in quintuplicates and HepG2 cells in quadruplicates were lysed in SDS lysis buffer (4 % SDS, 10 mM DTT in 50 mM Tris/HCl pH 7.5, cOmplete protease inhibitor cocktail), boiled at 95 °C for 8 min and sonicated for 15 min at 4 °C (Bioruptor). Alkylation of proteins was performed for 20 min in the dark by addition of 55 mM iodoacetamide in 50 mM Tris/HCl (pH 7.5), followed by normalization of the protein concentration (Pierce 660 nm Protein Assay). For each replicate, 50 µg proteins were precipitated with 4 volumes of pre-chilled acetone at −20 °C overnight, pelleted by centrifugation (15 000 x g, 10 min, 4 °C), washed with 80 % (v/v) acetone, air-dried and resuspended in 40 µl thiourea buffer (6 M urea, 2 M thiourea in 10 mM HEPES, pH 8.0). Proteins were pre-digested by addition of 1 µg LysC in 40 µl 50 mM ammonium bicarbonate (ABC) buffer (pH 8.0) for 3 hours at 25 °C. Digestion was completed with 1 µg Trypsin in 120 µl 50 mM ABC buffer (pH 8.0) for 16 hours at 25 °C. Following purification on C18 StageTips (HepG2 cells) or fractionation of peptides on SCX (7 fractions: flow-through, pH 11, pH 8, pH 6, pH 5, pH 4, pH 3) and purification on C18 StageTips (A549 cells) as described previously [82], peptides were loaded onto a 50 cm reverse-phase analytical column (75 µm column diameter; ReproSil-Pur C18-AQ 1.9 µm resin; Dr. Maisch) and separated using an EASY-nLC 1200 system (Thermo Fisher Scientific). A binary buffer system consisting of buffer A (0.1 % formic acid in H_2_O) and buffer B (80 % acetonitrile, 0.1 % formic acid in H_2_O) with a 120 min (A549 cells) gradient (5 to 30 % buffer B for 95 min, 30 to 95 % buffer B for 10 min, wash out at 95 % buffer B for 5 min, decreased to 5 % buffer B for 5 min, and 5 % buffer B for 5 min) was used at a flow rate of 300 nl/min. In contrast, peptides derived from HepG2 cells were separated with a 180 min gradient. Eluting peptides were directly analyzed on a Q-Exactive HF mass spectrometer (Thermo Fisher Scientific) operated in data-dependent acquisition mode with repeating cycles of one MS1 full scan (300-1 650 m/z, R=60 000 at 200 m/z) at an ion target of 3e6, followed by 15 MS2 scans of the highest abundant isolated and higher-energy collisional dissociation (HCD) fragmented peptide precursors (R=15 000 at 200 m/z). Peptide precursor isolation for MS2 scans was additionally limited by a maximum injection time of 25 ms and an ion target of 1e5 while repeated isolation of the same precursor was dynamically excluded for 20 s. Spectra were processed with MaxQuant (A549: version 1.6.6.0; HepG2: 1.6.10.43) using label-free quantification, match between runs, fixed carbamidometylation and variable N-acetylation and methionine oxidation. Peptides and proteins were identified by searching against the human proteome sequences (UniprotKB, 3.2016) controlled by a false discovery rate of 0.01 [83]. Further analyses were performed with Perseus (version 1.6.6.0) [84]. Proteins only identified by site, matching the reverse sequence, or annotated as contaminants were excluded from the analysis. Protein copy numbers were estimated using the Proteomic Ruler plugin for Perseus as previously described [85]. Briefly, raw protein intensities were separately averaged for each column and the histone proteomic ruler with a ploidy of three was applied as scaling mode with a total cellular protein concentration of 200 g/l. Quantification accuracy was further estimated using the standard plugin parameters.

### Mathematical modelling

Implementation of all mathematical models was performed in the form of ordinary differential equations in MATLAB 2017a. Parameter optimization and uncertainty analysis was performed using the data2dynamics framework for MATLAB [86] and model analysis was performed using the ode15s solver in MATLAB. Detailed information about the mathematical model and its underlying assumptions are reported in the Supplementary Material.

### Parameter estimation of the RIG-I core model

Total intracellular protein levels were gathered from a deep proteomic analysis of A549 and HepG2 cells (Supplementary Material). 5’ppp-dsRNA was assumed to be uniformly distributed post electro-transfection, corresponding to a cytoplasmic dsRNA concentration of 2 nM. No basal RIG-I signaling was accounted for, thus, yielding all phosphorylated protein levels to be zero prior to stimulation. Additionally, in the absence of any stimulus, no free IκBα was considered. Total protein levels were converted to concentrations using the reported cytoplasmic and nuclear volume of A549 cells (Vcyt=1.2e-12 L and Vnucl = 4.7e-13 L [87]), Nine kinetic rate constants were derived from literature while the remaining rate constants were optimized during the fitting process. The cytoplasmic degradation rate of 5’ppp-dsRNA was set to the fitted intracellular degradation rate constants of HCV RNA after electro-transfection [88]. Reported half-lives for RIG-I proteins in HepG2 cells were used to define the basal RIG-I degradation rate constant and the basal RIG-I synthesis rate constant was derived by utilizing *k*_syn_ = [RIG-I]_*t*=0_ · μ_RIG-I_ [89]. Degradation of IκBα and the binding rate constant to NF-κB by IκBα were taken from the work of Hoffmann et al. [90], expecting no strong differences in their reaction kinetics between A549 and human JURKAT cell lines. Due to the lack of experimental data regarding the temporal RIG-I activation and MAVS complex formation dynamics, reported values for k_RIG-I_, k_MAVS,_ and b_RIG-I_ were employed from an existing mathematical model of the virus-triggered type I IFN signaling pathway [26]. The degradation rate constant of IFN-β mRNA was further derived from its reported half-life in immortalized human bronchial epithelial cells [91].The eleven remaining rate constants were optimized using experimentally observed western blot intensities and IFN-β qPCR data. Therefore, experimentally observed, normalized intensity values were linked to the concentrations of model species with the help of scaling factors and background intensities (Supplementary Material). Background intensities for phosphorylated proteins were defined as mean of normalized intensity values for the first two time-points of the short-period experiment (0 min, 1 min). The remaining scaling factors were optimized during the optimization process alongside with the kinetic rate constants. Parameter fitting was performed by simultaneously minimizing the negative logarithm of the likelihood for the observed experimental data given our model parameters using a multi-start optimization method based on Latin Hypercube Sampling (LHS) as implemented in the data2dynamics framework for MATLAB [86,92]. The parameters were log10-transformed prior to optimization and a search space of at least six orders of magnitude was applied. To ensure the convergence towards a global minimum in the multi-dimensional parameter space, 1500 runs were performed using LHS-sampled initial parameter guesses. The sorted final goodness of fit for every optimization run is shown in the Supplementary Material and shows repeated convergence towards a global optimum.

### Parameter estimation of the coupled model of antiviral signaling

Whenever possible, parameter values were taken from the previously established RIG-I core model or the original IFN pathway model [24]. Already existing protein and mRNA species of the core model constitute the same initial values as reported previously and total protein levels of IFN pathway components were adjusted based on proteomics data. Basal levels of JAK/STAT model species were determined by simulating their steady state concentrations in absence of any stimulus (Supplementary Material). IFN and ISG levels were set to zero prior to stimulation.

To combine the RIG-I pathway model to the published JAK-STAT model, we included a production rate of IFNpre, i.e., the stimulus of JAK-STAT signaling, by the IFN-β mRNA species of our RIG-I pathway core model. The remaining structure of the JAK-STAT model was not changed, but merely additional ISG model species were introduced to better capture the antiviral state of a cell by our model. In total, we considered the expression of 6 genes in our model (C-C motif chemokine ligand 5 (CCL5), C-X-C motif chemokine ligand 10 (CXCL10), IFIT1, IFN-λ, RIG-I, MX1), in addition to the already existing upregulated species IκBα, IFNβ, IRF9 and SOCS. Expression of ISGs is under the regulation of phosphorylated IRF3 (CCL5, CXCL10, IFNλ) or under the regulation of both phosphorylated IRF and active ISGF3 (IFIT1, RIG-I, MX1). Depending on the availability of qPCR and western blot data, ISGs are included at the transcript level (CCL5, CXCL10), the protein level (RIG-I) or at both levels (IFIT1, IFN-λ, MX1) as species in our model. Furthermore, we account for a first order decay of mRNAs (denoted μ) and, when a protein species is included into the model, a translation and protein degradation process.

Only a single rate constant (k_68_) of the Maiwald model [24] was re-adjusted, thereby accounting for differences in the potential of secreted IFN to trigger JAK/STAT signaling in the experimental set-ups. All remaining reported rate constants were assumed to be identical in A549 and Huh7.5 cells). Among the newly introduced parameters, degradation rate constants were fixed based on reported half-lives whenever possible (µ_CCL5,mRNA_, µ_CXCL10,mRNA_, µ_IFN-λ,mRNA_, µ_MX1,mRNA_, µ_IFIT1,mRNA_, µ_MX1_, µ_IFIT1_) [93–100]. The degradation of IFN in the supernatant was assumed to be negligible and set to 0. Remaining kinetic rate constants were optimized based on experimentally observed levels of RIG-I protein, IFIT1 protein, IRF9 protein, phosphorylated STAT2 protein, IFIT1 mRNA, MX1 mRNA, IFNB1 mRNA levels, as well as IFN-α levels in the supernatant (Figure 5C). Model fitting was performed by minimizing the negative log likelihood using a multi-start approach based on Latin Hypercube sampling as implemented in the data2dynamics framework [86,92]. Moreover, scaling factors and background intensities were introduced to link experimentally observed intensities to intracellular concentrations. The resulting goodness of fit for every optimization run is shown in the Supplementary Material highlighting convergence towards a global optimum.

### Uncertainty analysis

Structural and practical identifiability of the core and coupled model were assessed by calculating the profile likelihood estimates for every optimized parameter as implemented in the data2dynamics framework [86,101]. Briefly, starting from the optimal set of parameters, every parameter was stepwise fixed at increasingly deviating values, while all other open parameters were re-adjusted. The relation between the goodness of fit for given changes in the parameter values allowed to identify parameter regions and likelihood-based 95 % confidence intervals [101]. The resulting likelihood profiles for the core model are shown in the Supplementary Material. The analysis revealed a linear dependency between the shared rate constant for the activation of TBK1/IKKε and the IRF3 phosphorylation rate constant by the two kinases. The shared rate constant for the activation of TBK1/IKKε was consequently fixed and the IRF3 phosphorylation rate was optimized. For all other fitted parameters of the RIG-I core model, no identifiability problems could be identified.

## Supporting information

Supplementary figured and tables

## Acknowledgments

We want to thank Vladimir G. Magalhães for the productive discussions and support, Maike Drechsler for excellent technical assistance, the DKFZ research group *Molecular Therapy of Virus-Associated Cancers* for IncuCyte-related support, Christoph Stein-Thöringer, Nyssa Cullin, and Rogier Gaiser for providing the MSD instrument and the associated technical support, Friedemann Weber and Georgios Panagiotidis (Giessen University) for experimental expertise, Konstantin Sparrer (Ulm University) for providing SARS-CoV2 ORF6, Volker Lohmann for the provision of cell lines, and Ralf Bartenschlager for providing an excellent research environment. We are grateful to the Helmholtz International Graduate School for Cancer Research for supporting SSB with a PhD stipend.

## Author contributions

This project was designed by SSB, DS, JF, JW, LK, and MB. Molecular cloning and live-cell imaging was carried out by SSB and data analyzed together with MB. Cell line generation was done by SSB and SW. Cell Culture experiments were conducted by SSB, JF, and CS and analyzed together with MB. Mass spectrometry experiments were performed and analyzed by CU and AP. Mathematical modeling was done by DS and LK. The manuscript was prepared by SSB and MB and edited and approved of by all authors. The work was supervised by MB and LK.

## Data and materials availability

Proteome data PRIDE-ID: PXD031406

