## Supplementary figured and tables for "High Resolution Kinetic Characterization and Dynamic Mathematical Modeling of the RIG-I Signaling Pathway and the Antiviral Responses"

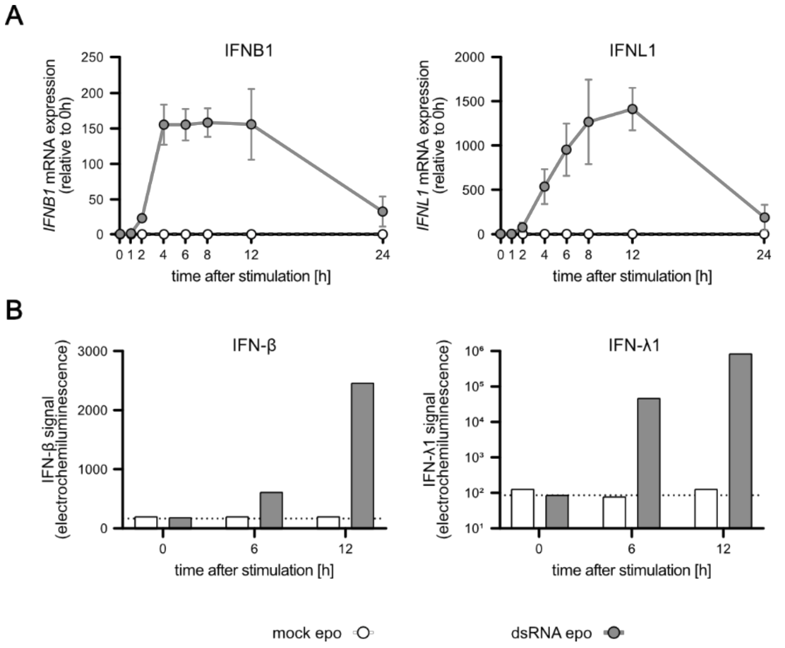


**Supplementary Figure S1. IFN mRNA expression and secretion upon synchronous stimulation in A549 wt cells.**

A549 wt cells were either mock electro-transfected or stimulated with 220 ng 5'ppp-dsRNA. **(A)** IFNB1 and IFNL1 mRNA expression was measured over time using qRT-PCR. Values were normalized to the housekeeping gene GAPDH and the 0 hour time point subsequently using $2^{-\Delta\Delta Ct}$. Graphs depict mean ± SD of three biologically independent experiments. **(B)** Secreted IFN-β and IFN-λ1 protein concentrations were determined using a multiplex immunoassay (U PLEX IFN Combo, Meso Scale Diagnostics) in mock or dsRNA stimulated A549 wt cells. Graphs depict electrochemiluminescent signals measured with the MESO QuickPlex SQ 120 instrument. Dashed lines indicate lower limit of detection.

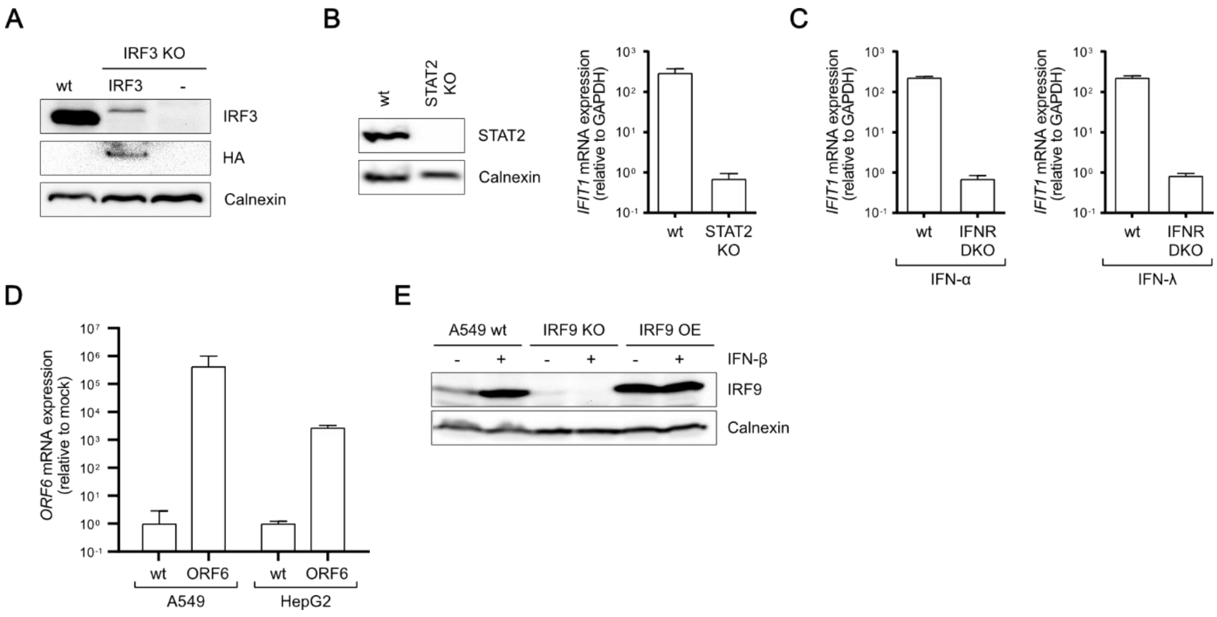


**Supplementary Figure S2. Expression control of KO or OE cell lines.**

**(A)** Western Blot analysis of basal IRF3 or IRF3-HA expression in A549 wt, A549 ROSA26-IRF3, and A549 IRF3 KO cells. **(B)** Western Blot analysis of basal STAT2 expression and IFIT1 mRNA expression upon stimulation with 100 IU/ml IFN-α in A549 wt and A549 STAT2 KO cells. **(C)** IFIT1 mRNA expression in A549 wt and A549 IFNR DKO cells upon IFN-α or IFN-λ stimulation for 16 hours. **(D)** ORF6 mRNA expression in A549 wt and HepG2 wt or ORF6 expressing cells. **(E)** IRF9 protein expression in A549 wt, A549 IRF9 KO, and A549 ROSA26-IRF9 (IRF9 OE) cells stimulated with or without 200 IU/ml IFN-β for 16 hours.

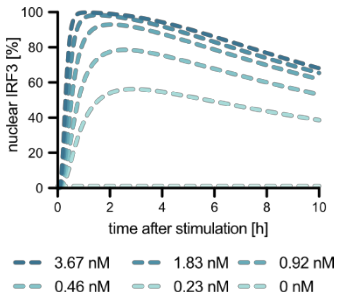


**Supplementary Figure S3:** Predicted nuclear translocation of pIRF3 upon electroporation of varying dsRNA levels.

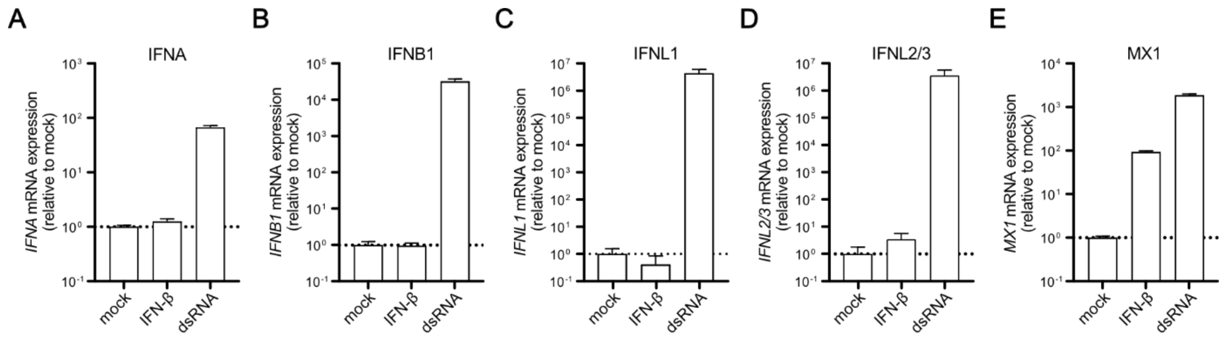


**Supplementary Figure S4. IFN and MX1 mRNA expression upon IFN and dsRNA stimulation in A549 wt cells.**

A549 wt cells were either mock stimulated, stimulated with 200 IU/ml IFN-β or transfected with 100 ng dsRNA using Lipofectamine 2000 for 16 hours. **(A)** IFNA, **(B)** IFNB1, **(C)** IFNL1, **(D)** IFNL2/3, as well as **(E)** MX1 mRNA expression was analyzed using qRT-PCR. Values were normalized to the housekeeping gene GAPDH and the mock stimulation subsequently using $2^{-\Delta\Delta Ct}$. Dashed lines indicate mock control levels.

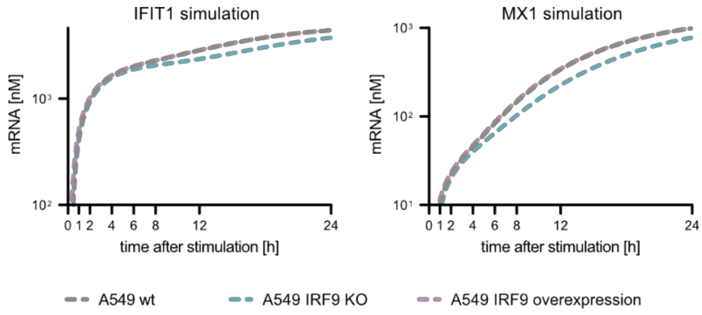


**Supplementary Figure S5. Model simulations with increased or decreased IRF9 levels.**

IRF9 levels were increased (10-fold, blue line) or knocked-out (red line) and IFIT1 and MX1 mRNA expression was simulated and compared to wt (black line) IRF9 levels.

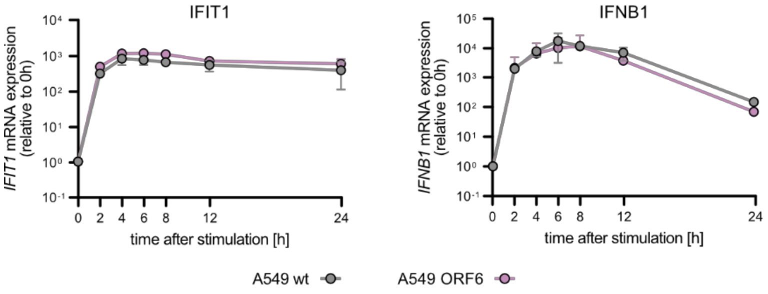


**Supplementary Figure S6. IFIT1 and IFNB1 mRNA expression upon SARS-CoV-2 ORF6 expression in A549.**
A549 wt or A549 cells stably expressing SARS-CoV-2 ORF6 were synchronously stimulated with 220 ng 5'ppp-dsRNA. IFIT1 and IFNB1 mRNA expression was monitored over time. Values were normalized to the housekeeping gene GAPDH and the 0 hour time point subsequently using $2^{-\Delta\Delta Ct}$. Graphs depict mean ± SD of two biologically independent experiments.

**Supplementary Table S1.** Guide RNAs for CRISPR/Cas9 KO generation.

| **Gene** | **Sequence (5' – 3')** |
|---|---|
| IFNR DKO (double KO of IFNAR1 and IFNLR1) | gaccctagtgctcgtcgccg<br>caccggagtaccagatcatgccac |
| STAT2 | gtcgaatgtccacaggcagg |

**Supplementary Table S2.** Primers for qRT-PCR.

| **Gene** | **Sequence (5' – 3')** |
|---|---|
| GAPDH | tcggagtcaacggatttggt<br>ttcccgttctcagccttgac |
| IFIT1 | gaatagccagatctcagaggagc<br>ccatttgtactcatggttgctgt |
| IFNA | agccatctctgtcctccatgag<br>gatctcatgatttctgctctga |
| IFNB1 | cgccgcattgaccatcta<br>gacattagccaggaggttctc |
| IFNL1 (IL-29) | ggtgactttggtgctaggct<br>tgagtgactcttccaaggcg |
| IFNL2/3 (IL-28) | ctgccacatagcccagttca<br>agcgactcttctaaggcatct |
| TNFAIP3 | tcctcaggctttgtatttgagc<br>tgtgtatcggtgcatggtttta |
| MX1 | accattccaaggaggtgcag<br>tgcgatgtccacttcggaaa |

| | |
|---|---|
| RIG-I | ccctggtttagggaggaaga<br>tcccaactttcaatggcttc |
| CCL5 | gctgtcatcctcattgctactg<br>tggtgtagaaatactccttgatgtg |
| ISG15 | acagccatgggctggga<br>ccttcagctctgacaccgac |
| ORF6 | atgtttcatctcgttgactttcagg<br>ttaatcaatctccattggttgctctt |
